# Maternal Western-style Diet Promotes Immune Tolerance and Liver Sinusoidal Endothelial Cell Dysfunction in Nonhuman Primate Juvenile Offspring Liver

**DOI:** 10.64898/2026.01.06.697995

**Authors:** Michael J. Nash, Benjamin N. Nelson, Molly McGuckin, Rachel C. Janssen, Kenneth L. Jones, Saif I. Al-Juboori, Evgenia Dobrinskikh, Paul Kievit, Kjersti M. Aagaard, Carrie E. McCurdy, Maureen Gannon, Stephanie R. Wesolowski, Jacob E. Friedman

**Affiliations:** Department of Pediatrics, University of Colorado Anschutz, Aurora, CO, USA; Harold Hamm Diabetes Center, University of Oklahoma Health Campus, Oklahoma City, OK, USA; Bioinformatic Solutions, Sheridan, WY, USA; Division of Cardiometabolic Health, Oregon Health Science University, Oregon National Primate Research Center, Beaverton, OR, USA; Division of Neuroscience, Oregon Health Science University, Oregon National Primate Research Center, Beaverton, OR, USA; HCA Healthcare and HCA Healthcare Research Institute, Nashville, TN and HCA Healthcare Texas Maternal Fetal Medicine, Houston, TX, USA; Fetal Care and Surgery Center, Division of Fetal Medicine and Surgery, Boston Children’s Hospital, Harvard Medical School, Boston, MA, USA; Department of Human Physiology, University of Oregon, Eugene, OR, USA; Department of Medicine, Division of Diabetes, Endocrinology, and Metabolism, Vanderbilt University Medical Center, Nashville, TN, USA; Department of Biochemistry and Physiology, University of Oklahoma Health Campus, Oklahoma City, OK, USA; Present address: Department of Medicine, David Geffen School of Medicine, University of California Los Angeles, Los Angeles, California, USA

## Abstract

Maternal Western-style diet (mWSD) consumption during pregnancy and lactation is associated with developmental programming of metabolic dysfunction-associated steatotic liver disease (MASLD) in offspring. To understand the roles of immune and endothelial cells, we used single-cell RNA-sequencing of liver non-parenchymal cells from 3-year-old juvenile nonhuman primates exposed to mWSD during their gestation through weaning, followed by control diet consumption after weaning. We identified unique clusters of macrophages in mWSD-exposed juvenile livers with non-reparative, pro-fibrotic phenotypes characterized by predicted inactivation of NF-κB, decreased oxidative phosphorylation, and gene expression facilitating hepatic stellate cell-macrophage interactions. Kupffer cell and dendritic cell (DC) numbers were decreased by mWSD exposure, with inactivation of inflammatory and antigen presentation pathways in DCs, supporting DC immaturity. B cells increased in mWSD-exposed offspring, with RNA showing reduced inflammation and impaired differentiation, while T cells had RNA profiles consistent with apoptosis and reduced inflammatory function. mWSD exposure increased clusters of liver sinusoidal endothelial cells (LSECs), with activation of inflammation and proliferation pathways but decreased immune cell communication. Immunocytochemistry and RNAscope identified increased association of LSECs and immune cells in periportal regions. In summary, mWSD exposure during gestation and lactation selectively modulated LSEC-immune cell interactions consistent with immune tolerant B and T cells and fibrogenic pathways together with decreased pro-resolving macrophages in juvenile offspring. We conclude that mWSD exposure establishes an immune-tolerant environment in offspring liver, marked by a durable, pro-fibrotic microenvironment that resists postnatal dietary correction.

## Introduction

Pediatric MASLD affects approximately 10–34% of children and adolescents globally^1^. A higher prevalence is observed among children born to mothers with obesity or poor dietary habits^2,3,4^, supporting that it can begin in utero. Pediatric MASLD is characterized by variable degrees of steatosis, inflammation, and periportal fibrosis, predominantly in the periportal (zone 1) region, that may not always follow in succession^5,6^. In contrast, adult MASLD is characterized by successive steatosis, inflammation, and fibrosis, with fibrosis developing first in pericentral and perisinusoidal regions^7^. Multiple developmental studies in mice^8,9^, nonhuman primates (NHP)^10,11^, and humans^12,13^ indicate that exposure to maternal Western-style diet (mWSD) or maternal obesity increases the risk and severity of MASLD in offspring^4^. However, the mechanisms that drive liver fibrosis early in pediatric MASLD remain poorly understood.

Hepatic inflammation drives hepatic stellate cell (HSC) activation and the progression of fibrosis in adult MASLD^7^. A balance between inflammation and the resolution of inflammation is important for liver function and tissue repair^14^. While chronic low-grade inflammation is associated with hepatocellular injury and fibrosis in MASH^15^, restoration of tissue function and liver regeneration is dependent on both an acute inflammatory response and subsequent action of reparative, non-inflammatory tissue resident macrophages, as well as recruited bone marrow-derived macrophages that facilitate hepatocyte regeneration and clearance of apoptotic cells^16,17^. Liver sinusoidal endothelial cells (LSECs) control macrophage phenotypes in a niche-dependent manner^18^ by informing the innate and adaptive immune systems^19^. Dysfunctional LSECs release inflammatory mediators, leading to the recruitment of inflammatory cells and subsequent liver injury and inflammation and parallels MASLD severity^20^. However, little is known about how mWSD exposure regulates LSECs or immune cell functions and initiates pediatric MASLD risk in exposed offspring.

Single-cell resolution RNA-sequencing (scRNA-seq) in adult humans with MASLD has identified pathological mechanisms of liver fibrosis^21^ and a role for both adaptive and innate immune cells^22^ and LSECs^23^, supporting the importance of cell-cell interactions in the liver of adult MASLD patients. However, the earlier onset and differences in MASLD presentation in children compared with adults suggests the presence of unique drivers and disease mechanisms. Exposure to mWSD in utero, representing diet composition and associated maternal and placental responses, may initiate liver injury early in development^4^. More specifically, crosstalk between LSECs and innate and adaptive immune cells may be critical mediators of immune tolerance. Immune cell tolerance can lock the liver into a fibrotic state by creating a suppressive microenvironment in which regulatory T cells and immunosuppressive macrophages restrain inflammation but also inhibit the activation of pro-resolution macrophages and cytotoxic cells that could clear scar-forming cells^24,25^ thus creating a maladaptive response to liver damage. Such mechanisms are classically observed in liver cancer^26,27^ and hepatitis C^28^.

Here, we used scRNA-seq combined with histological and metabolomic analysis in our well-characterized NHP model of mWSD exposure that demonstrates persistent portal fibrosis in fetal and juvenile offspring^10,29,30,31^. We show the molecular and cellular consequences of mWSD exposure on postnatal hepatic immune and endothelial cell phenotypes in 3-year-old juvenile offspring. Our results demonstrate that periportal zones in mWSD-exposed offspring liver are home to a pro-fibrotic niche with immune-tolerant immune cells despite switching to a chow diet at weaning. This finding supports that developmental remodeling of innate and adaptive immunity toward a novel immune tolerant phenotype contributes to persistent, non-reparative periportal fibrosis in our NHP model of pediatric MASLD, with in utero origins.

## Results

### Phenotypic characterization of intrahepatic non-parenchymal mononuclear cells with scRNA-seq

To identify how mWSD exposure disrupts hepatocellular development in exposed offspring, we obtained scRNA-seq profiles from liver non-parenchymal mononuclear cells from a subset of 3-year-old offspring exposed to mWSD during pregnancy and lactation or maternal chow diet (mCD) (Fig. 1A; Supplementary Table 1) from our NHP model^10^. We identified populations of T cells, B cells, Kupffer cells (KCs; resident macrophages), other (non-KC) macrophages, dendritic cells (DCs), endothelial cells, cholangiocytes, and stellate cells (Fig. 1B; Supplementary Fig. 1). These cell populations are similar to what was observed in adult human liver scRNA-seq^22,32^. The small populations of stellate cells and hepatocytes were not analyzed further, as they represent carryover from our liver cell isolation protocol^33^, which was designed to enrich for mononuclear immune and endothelial cells. Results for cholangiocytes have been reported^30^. Numbers of differentially expressed genes (DEGs) in mWSD compared with mCD groups per cell population ranged from 302 to 842 (Fig. 1C). The different cell populations observed across samples are shown in Fig. 1D. In mWSD- relative to mCD-exposed offspring livers, the proportions of B cells and endothelial cells were increased (proportion >0.7), while cholangiocytes, DCs, and KCs were relatively decreased (proportion <0.35), and macrophages and T cells were slightly decreased (proportion <0.5, Fig. 1E; Supplementary Table 2). In the following sections, we further interrogated these cell populations and used Ingenuity Pathway Analysis (IPA), combined with validation of select targets in select immune and endothelial cell populations, to identify predicted functional roles of DEGs regulated by mWSD exposure.

**Figure 1.**
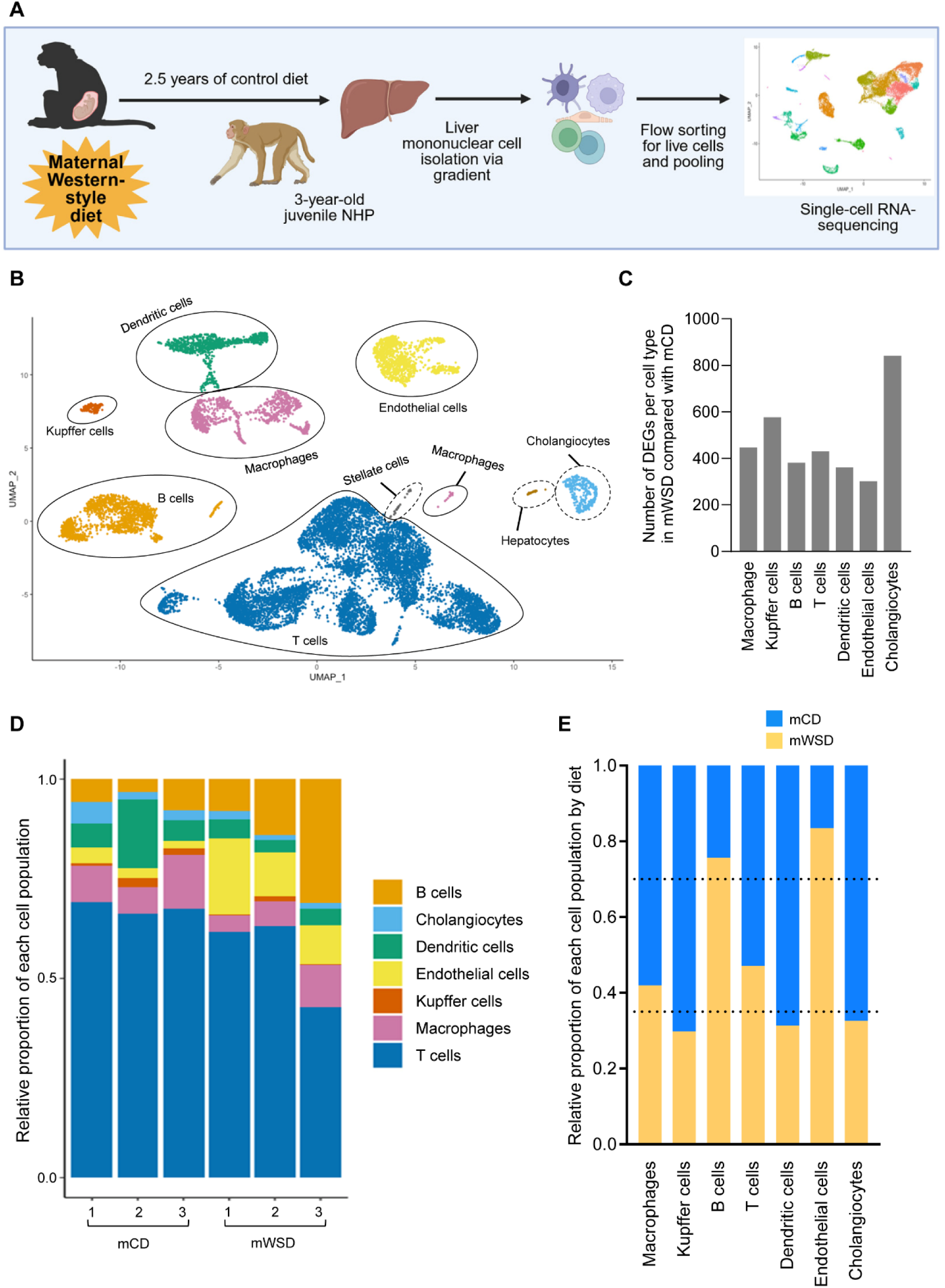
Identification of intrahepatic mononuclear cells with scRNA-seq analysis. **A** Nonhuman primate (NHP) model of mWSD exposure and experimental design for scRNA-seq of offspring liver mononuclear cells. Created in BioRender. **B** UMAP showing cell populations identified in both mCD- and mWSD-exposed livers. Cell clusters with dashed outline were not analyzed herein. **C** Number of differentially expressed genes (DEGs) between mCD- and mWSD-exposed livers per cell population. **D** Proportion of each cell population identified per individual NHP sample. **E** Relative proportion of each cell population by maternal diet. Dashed lines represent boundaries to show decreased (lower line) and increased (upper line) proportions in mWSD- vs. mCD-exposed liver. n = 3 mCD, n = 3 mWSD.

### mWSD exposure is associated with decreased inflammatory phenotypes across clusters of liver macrophages

Macrophage activation promotes MASLD progression in adults^34^. Unexpectedly, in mWSD-exposed livers, the DEGs in the non-KC macrophage populations predicted inactivation of pro-inflammatory pathways compared with mCD-exposed livers, including macrophage classical activation, interferon gamma, IL-1 and IL-6, and the anti-inflammatory IL-10 pathway (Fig. 2A; Supplementary Table 3). Conversely, these macrophages also had predicted activation of pathways for interferon alpha/beta, interferon signaling, and death receptor signaling, all consistent with apoptosis, as well as predicted activation of PPAR and complement cascade (Fig. 2A). Expression of individual pro-inflammatory genes such as *IL1B* and *NFKBIA* and mitochondrial-encoded electron transport genes including *COX1*, *COX2*, *COX3*, and *ND4* were decreased in macrophages from mWSD-exposed livers (Fig. 2B). Genes mediating stellate cell activation and interaction with macrophages, including *MSRB1* and *TIMP1*, were increased, as were genes controlling complement signaling including *CFD* and *CFP* (Fig. 2B; Supplementary Table 4).

**Figure 2.**
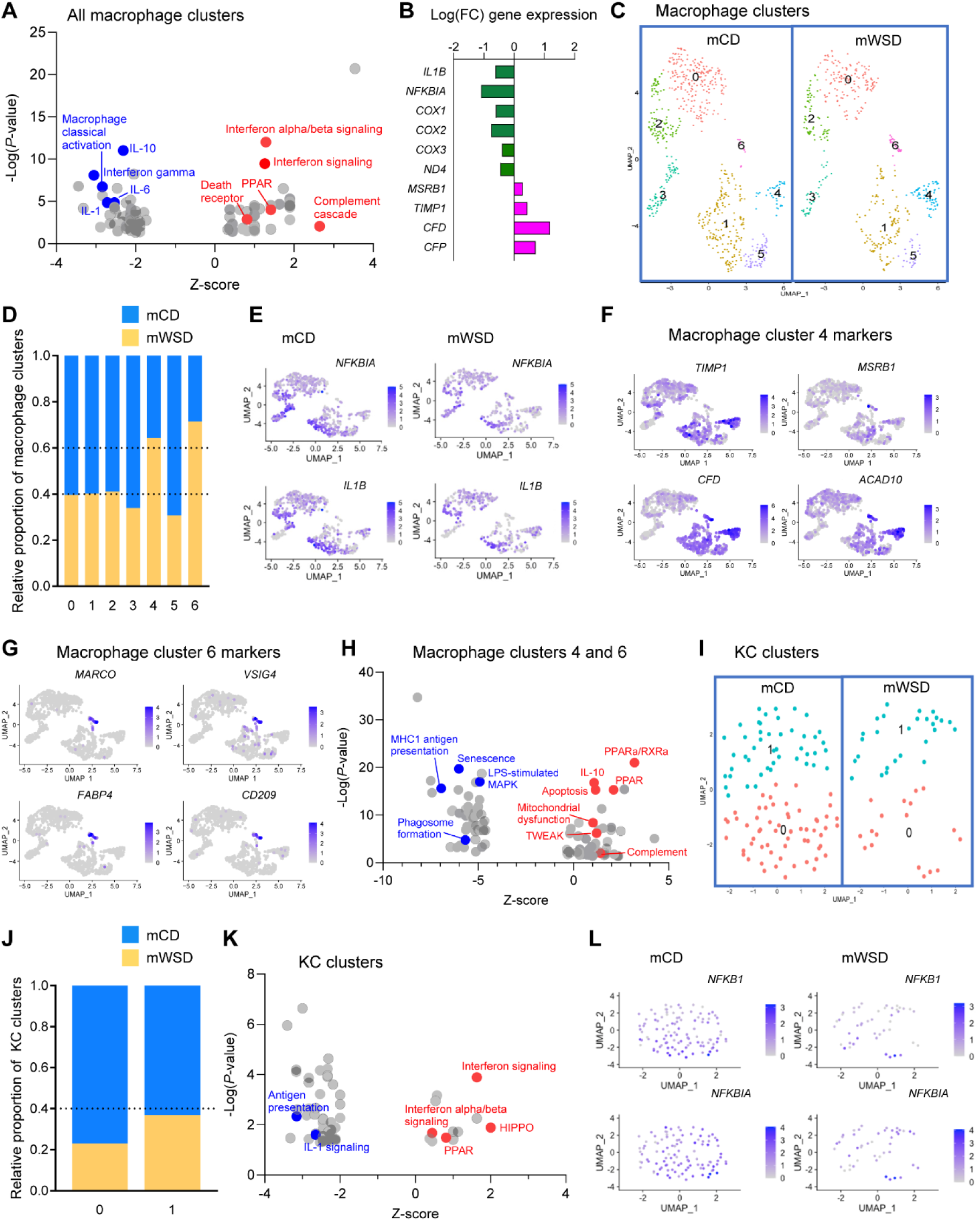
mWSD exposure is associated with decreased inflammatory phenotypes in hepatic macrophages. **A** Activation Z-scores and *P*-values for the significant top 50 inactivated and all activated (46) pathways (by *P*-value) predicted from DEGs comparing mWSD- vs. mCD-exposed non-KC macrophages. **B** Fold-change in expression of select DEGs in mWSD- vs. mCD-exposed macrophages. **C** UMAP of non-KC macrophage population with clusters numbered 0-6 identified in mCD and mWSD groups. **D** Relative proportion of each macrophage cluster by diet. Dashed lines represent boundaries to show decreased (lower line) and increased (upper line) proportions in mWSD- vs. mCD-exposed macrophages. **E** UMAP of non-KC macrophage populations showing expression of inflammatory *NFKBIA* and *IL1B* genes in all mCD- and mWSD-exposed macrophage clusters. UMAP of non-KC macrophages showing the distribution of four genes defining cluster 4 (**F**) and cluster 6 (**G**) in mCD and mWSD groups combined. **H** Activation Z-scores and *P*-values for the significant top 50 inactivated and top 50 activated pathways (by *P*-value) predicted by DEGs in macrophage clusters 4 and 6. **I** UMAP of Kupffer cell (KC) populations split by maternal diet and cluster, numbered 0-1. **J** Relative proportion of each KC cluster by diet. Dashed line represents a boundary to show decreased proportion in mWSD- vs. mCD-exposed KCs. **K** Activation Z-scores and *P*-values for the significant top 50 inactivated and all activated (13) pathways (by *P*-value) predicted from DEGs in KCs. **L** UMAP of KCs showing distribution of expression of pro-inflammatory *NFKB1* and *NFKBIA* by maternal diet. **A**, **H**, **K** Select red pathways indicate positive Z-score and activation; blue pathways, negative Z-score and inactivation; grey, top activated or inactivated pathways not highlighted. **E**-**G**, **L** Light purple corresponds to lower expression of select gene; dark purple, higher expression.

Within the macrophage population, we identified seven unique clusters (Fig. 2C). In mWSD-exposed livers, clusters 4 and 6 were overrepresented (proportion >60), while the other clusters were decreased (proportion <40) compared with mCD-exposed livers (Fig. 2D; Supplementary Table 2). Importantly, *NFKBIA* and *IL1B* were two DEGs with decreased expression across most macrophage clusters in mWSD-exposed livers (Fig. 2E). Notably, regardless of maternal diet, cluster 4 expressed high levels of *TIMP1* and *MSRB1*, as well as *CFD*, a major complement cascade factor, and *ACAD10*, which controls fatty acid oxidation (Fig. 2F). Cluster 6 expressed high levels of *MARCO* and *VSIG4*, genes shown to mark portal-associated and tolerogenic macrophages in humans^32^, *FABP4*, associated with M1-mediated inflammation^35^, and *CD209*, associated with an immunotolerant macrophage phenotype^36^ (Fig. 2G). Given the roles of these defining genes in clusters 4 and 6 and the overrepresentation of these clusters in mWSD-exposed livers, we focused on the DEGs in clusters 4 and 6. This analysis showed predicted inactivation of pathways for senescence, phagosome formation, MHC class I antigen presentation, and lipopolysaccharide (LPS)-stimulated MAPK signaling and activation of pathways for PPARa/RXRa, IL-10, PPAR, mitochondrial dysfunction, apoptosis, complement signaling, and TWEAK pro-fibrotic signaling, indicating that macrophages from mWSD-exposed livers are oriented toward immune tolerance and anti-inflammatory behavior rather than inflammation and tissue homeostasis (Fig. 2H; Supplementary Table 5).

### KCs in mWSD-exposed livers are lower but have similar non-inflammatory phenotypes as other intrahepatic macrophages

KCs have “restorative” functions in resolving fibrosis in MASH in adults^37^. The overall KC population was lower in mWSD-exposed livers (Fig. 2I) with the two clusters representing <40% compared with mCD-exposed livers (Fig. 2J; Supplementary Table 2). Among the DEGs regulated by mWSD exposure within both clusters, predicted inactivated pathways included antigen presentation by MHC class 1 and IL-1 signaling, and predicted activated pathways included interferon signaling, interferon alpha/beta signaling, and PPAR (Fig. 2K; Supplementary Table 6), similar to other macrophages from mWSD-exposed livers. A predicted activated pathway unique to KCs included the pro-fibrotic HIPPO pathway (Fig. 2K). Across KC clusters, patterns of DEGs by diet were relatively uniform and pro-inflammatory genes *NFKB1* and *NFKBIA* were decreased in mWSD-exposed livers (Fig. 2l). Overall, KCs from mWSD-exposed livers had a similar phenotype as non-KC macrophages, consistent with a non-reparative, immune tolerant phenotype.

### mWSD-exposed livers have evidence of reduced intrahepatic B cell activation and differentiation

Given the macrophage phenotype supporting impaired function and antigen presentation, we next investigated the adaptive immune cells. Seven unique clusters of B cells were observed (Fig. 3A), with cluster 6 representing a plasma cell phenotype (expressing *JCHAIN*, *IGKC*, *IGHM*; Supplementary Fig. 2). There was an overall increase in intrahepatic B cells with mWSD exposure, with cluster 4 being almost nonexistent in mCD-exposed livers (proportion <0.05 in mCD compared with >0.95 in mWSD; Fig. 3B; Supplementary Table 2). Functional annotation of DEGs by mWSD exposure in total B cell populations, across all clusters, predicted inactivation of TNF and IL-3 inflammatory pathways that potentiate antibody production and B cell differentiation^38^, JAK/STAT signaling, oxidative phosphorylation, and B cell signaling (Fig. 3C; Supplementary Table 7). B cells from mWSD-exposed offspring also had a predicted activation of pathways including interferon alpha/beta signaling, interferon signaling, mitochondrial dysfunction, PPAR, and PPARa/RXRa signaling (Fig. 3C). Interestingly, the DEGs in highly enriched cluster 4 in mWSD-exposed offspring predicted inactivated inflammatory TNF and IL-1 signaling, oxidative phosphorylation, antioxidant signaling (KEAP1/NFE2L2), MHC1 antigen presentation, and B cell receptor signaling (Fig. 3D; Supplementary Table 8). Conversely, predicted activated pathways in cluster 4 included mitochondrial dysfunction, anti-inflammatory IL-10 signaling, and VEGF pathway, suggesting endothelial crosstalk (Fig. 3D). Thus, in addition to an overall increase in infiltration of B cells in mWSD-exposed livers, the predicted pathways were skewed toward a distinct non-inflammatory B cell phenotype with reduced antigen presentation.

**Figure 3.**
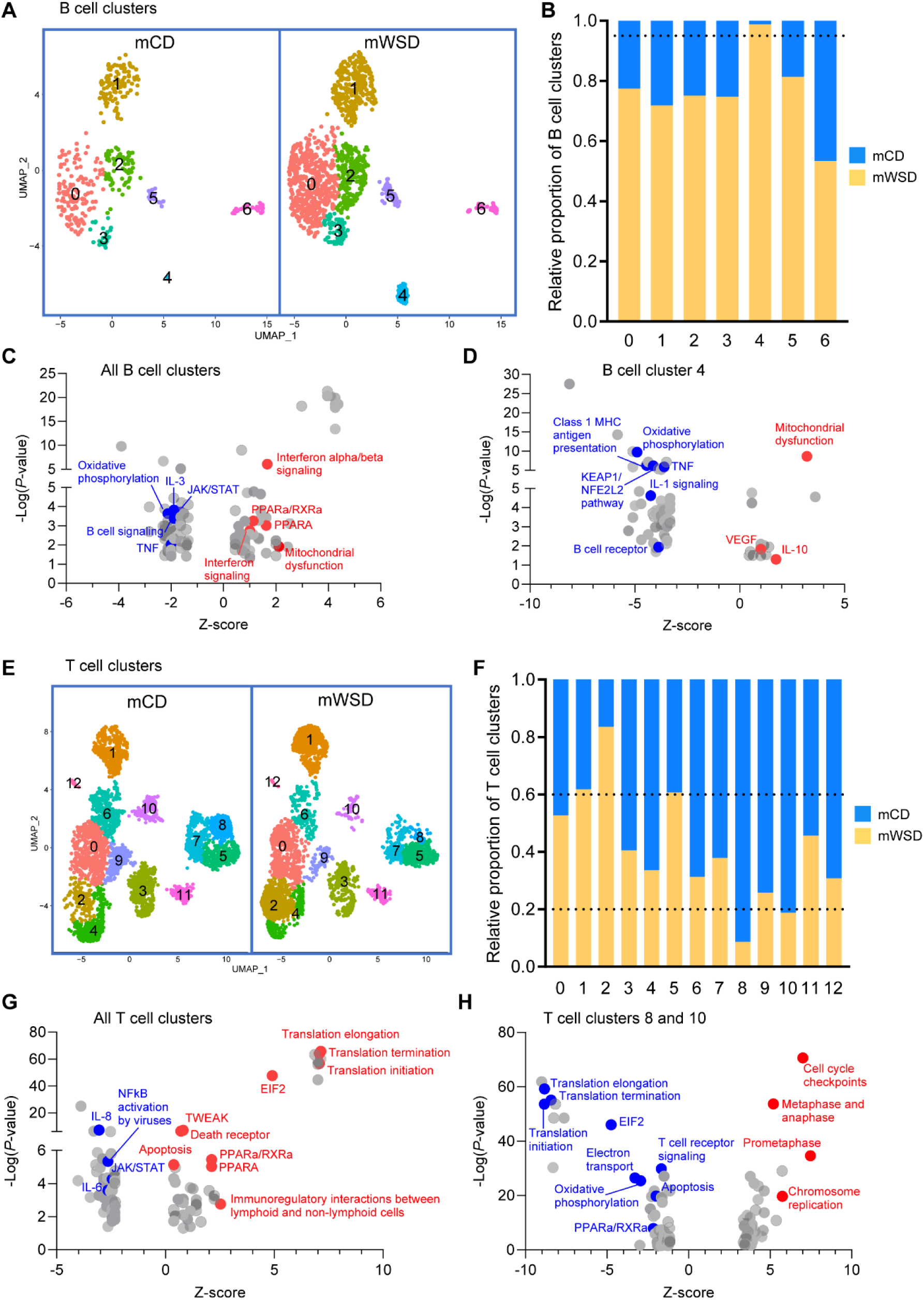
mWSD-exposed livers have evidence of reduced B cell activation and differentiation and an absence of inflammatory signatures in T cells. **A** UMAP of B cells split by maternal diet showing clusters numbered 0-6. **B** Relative proportion of each B cell cluster in mCD and mWSD groups. Dashed line represents a boundary to show increased proportion in mWSD- vs. mCD-exposed B cells. **C** Activation Z-scores and *P*-values for the significant top 50 inactivated and top 50 activated pathways (by *P*-value) derived from comparison of mWSD- vs. mCD-exposed total B cells. **D** Activation Z-scores and *P*-values for the significant top 50 inactivated and all activated (15) pathways (by *P*-value) derived from markers of B cell cluster 4. **E** UMAP of T cells split by maternal diet showing clusters numbered 0-12. **F** Relative proportion of each T cell cluster in mCD- and mWSD-exposed livers. Dashed lines represent boundaries to show decreased (lower line) and increased (upper line) proportions in mWSD- vs. mCD-exposed T cells. **G** Activation Z-scores for the significant top 50 inactivated and all activated (42) pathways (by *P*-value) derived from comparison of mWSD- vs. mCD-exposed total T cells. **H** Activation Z-scores for the significant top 50 inactivated and top 50 activated pathways (by *P*-value) derived from genes defining T cell clusters underrepresented in mWSD-exposed livers (clusters 8 and 10). **C**-**D**, **G**-**H** Select red pathways indicate positive Z-score and activation; blue pathways, negative Z-score and inactivation; grey, top activated or inactivated pathways not highlighted.

### mWSD-exposed intrahepatic T cells have a tolerant-like phenotype with lack of inflammatory signatures

In the T cell population, we identified 13 distinct clusters (Fig. 3E). Expression of CD4 and CD8 are similar in T cells from mCD- and mWSD-exposed livers (Supplementary Fig. 3). Eight out of 13 clusters identified were decreased in mWSD-exposed livers compared with mCD-exposed livers, although clusters 1, 2, and 5 were relatively increased (proportion >0.6, Fig. 3F; Supplementary Table 2). Across all T cell clusters, predicted inactivated pathways in mWSD-exposed offspring included NF-κB activation by viruses, IL-6, IL-8, and JAK/STAT, supporting downregulation of classical T helper inflammation (Fig. 3G; Supplementary Table 9). Predicted activated pathways included protein synthesis (EIF2, translation initiation, elongation, and termination), metabolism (PPAR, PPARa/RXRa), apoptosis and death receptor signaling, immunoregulatory interactions between lymphoid and non-lymphoid cells, and TWEAK (Fig. 3G). Predicted inactivated pathways in clusters 8 and 10 (underrepresented in mWSD-exposed livers, proportion <0.2, Fig 3F) included translation (translation initiation, elongation, termination, EIF2), oxidative phosphorylation, electron transport, PPARa/RXRa, apoptosis, and T-cell receptor signaling (Fig. 3H; Supplementary Table 10). Predicted activated pathways included cell cycle checkpoints, anaphase and metaphase, prometaphase, and chromosome replication (Fig. 3H). Thus, expression profiles in T cell populations from mWSD-exposed livers predicted reduced T cell division and inactivation of pathways regulating effector function, suggesting reduced T helper activity and increased immune tolerance. Notably, one of the hallmarks of early T cell tolerance is a decrease in protein translation, a regulatory checkpoint early in T cell activation that helps enforce self-tolerance^39^.

### mWSD exposure reduces activation of DCs in livers and shifts both the innate and adaptive immune system toward a non-inflammatory phenotype

DCs have a key role in antigen presentation, immune cell activation, and immune tolerance^40^. Four clusters of intrahepatic DCs were identified in juvenile NHP livers (Fig. 4A). The proportions of DC clusters were decreased in mWSD-exposed livers compared with mCD-exposed livers (proportion <0.4, Fig. 4B; Supplementary Table 2). Phagocytosis, a pathway key for DC activation, was predicted to be inactivated in DCs from mWSD-exposed livers, whereas oxidative stress-mediated senescence, PPARa/RXRa, which triggers DC apoptosis^41^, IL-1B, immunoregulatory interactions between lymphoid and non-lymphoid cells, and immunogenic cell death were predicted to be activated (Fig. 4C; Supplementary Table 11). Thus, RNA profiles in DCs from mWSD-exposed livers suggest impaired DC maturation, antigen presentation, and inflammation, with increased immune tolerance.

**Figure 4.**
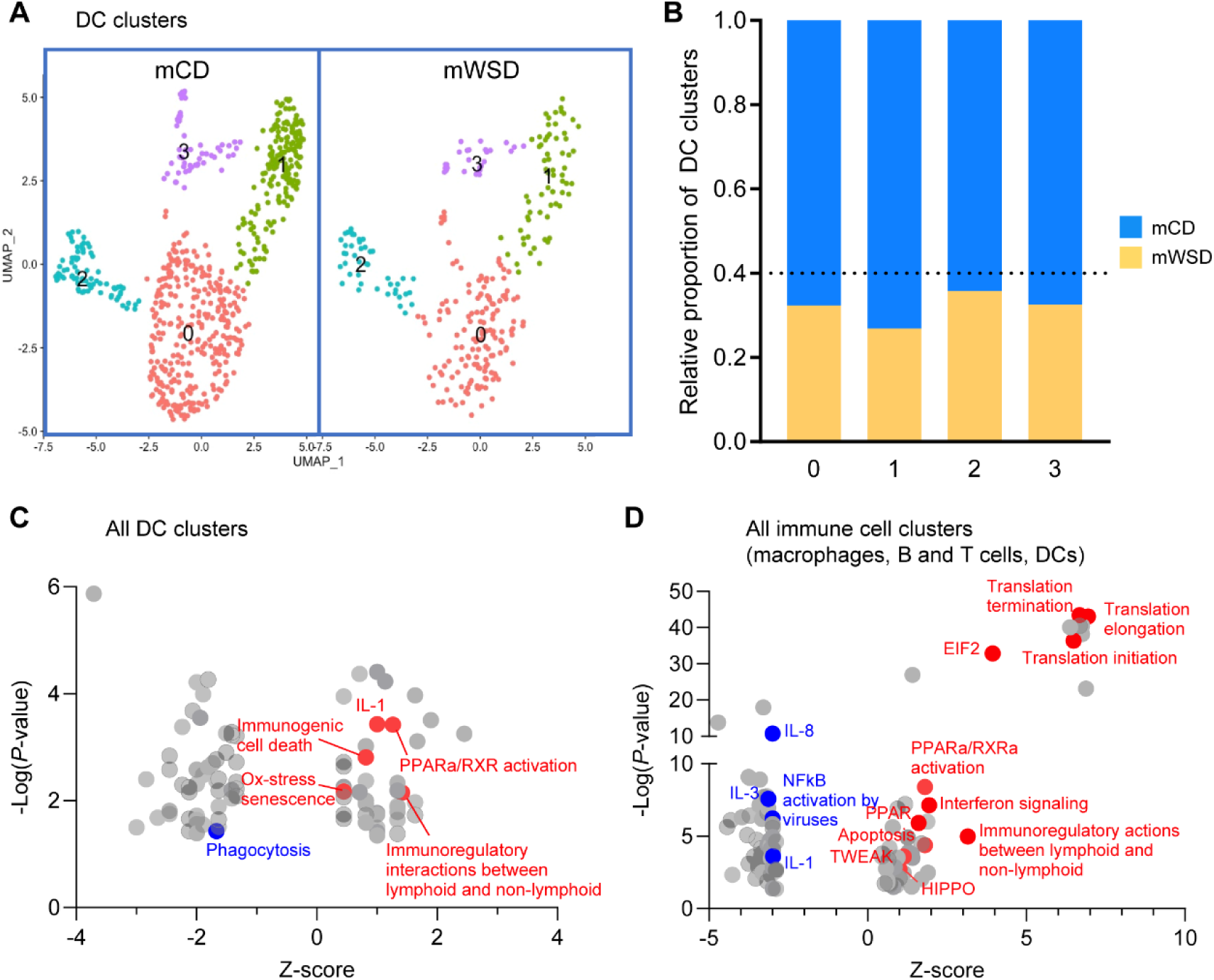
mWSD exposure reduces activation of hepatic DCs and shifts the innate and adaptive immune system toward a non-inflammatory phenotype. **A** UMAP of dendritic cells (DC) split by maternal diet showing clusters numbered 0-3. **B** Relative proportion of each DC cluster in mCD and mWSD groups. Dashed line represents a boundary to show decreased proportions in mWSD- vs. mCD-exposed DCs. **C** Activation Z scores and *P*-values for the significant top 50 inactivated and all activated (41) pathways (by *P*-value) derived from comparison of mWSD- vs. mCD-exposed total DCs. **D** Activation Z scores and *P*-values for the significant top 50 inactivated and top 50 activated pathways (by *P*-value) derived from comparison of mWSD- vs. mCD-exposed total immune cells (macrophage, KCs, DCs, B cells, T cells). **C**-**D** Select red pathways indicate positive Z score and activation; blue pathways, negative Z-score and inactivation; grey, top activated or inactivated pathways not highlighted.

Given the mWSD exposure-driven predicted remodeling of both innate and adaptive immune cell phenotypes with evidence of DC dysfunction, we analyzed all liver immune cells (macrophages, B cells, T cells, and DCs) together to identify phenotypes shared among DEGs. Predicted inactivated pathways included IL-1, IL-3, and IL-8 signaling and NF-κB activation by viruses (Fig. 4D; Supplementary Table 12). Predicted activated pathways included various protein translation processes and EIF2 signaling, interferon signaling, PPAR, PPARa/RXRa, apoptosis, immunoregulatory actions between lymphoid and non-lymphoid cells, and TWEAK and HIPPO signaling, both of which are implicated in fibrosis and HSC activation^42,43^ (Fig. 4D). This suggested a shared immunotolerant, non-inflammatory, pro-fibrotic phenotype across immune cell clusters in mWSD-exposed livers.

### mWSD exposure is associated with expansion of LSECs and activation of oxidative stress and proliferation pathways

LSECs regulate hepatic inflammation, fibrosis, and regression of fibrosis through intrahepatic immune cell tolerance^44,45^. We identified six clusters of intrahepatic endothelial cells (Fig. 5A) present in both mWSD- and mCD-exposed livers. Although most clusters increased with mWSD exposure (proportion >0.8), cluster 2 was similar to mCD-exposed livers and cluster 5 was underrepresented in mWSD-exposed livers (proportion <0.25, Fig. 5B; Supplementary Table 2). Importantly, endothelial cell clusters overrepresented in mWSD-exposed livers expressed *LYVE1*, indicating lymphatic or LSEC origin^46^. However, clusters 2 and 5 were defined by expression of *VWF*, indicating vascular endothelial cells (VECs) of blood vessel origin^47^ (Fig. 5C). Importantly, *PROX1*, used as a marker for lymphatic cells^46^, was expressed in only a very small proportion of the endothelial cells (Supplementary Fig. 4). Thus, the majority of endothelial cells in mWSD-exposed livers are LSECs based on expression markers. In contrast to what we observed in immune cells, expression of inflammatory genes such as *NFKBIA* was increased by mWSD exposure across nearly all endothelial cell clusters, a pattern shared with *TGFB1*, a marker of endothelial activation (Fig. 5D). Across all endothelial cell clusters, predicted inactivated pathways included mitochondrial dysfunction and macrophage classical activation (Fig. 5E; Supplementary Table 13). Predicted activated pathways included adaptive immune cell recruitment and stimulation (B cell receptor signaling, MHC class II antigen presentation), inflammatory interferon alpha/beta and PPAR, mitotic metaphase and anaphase, prometaphase and G2-G2/M phases, and pathways driving endothelial activation and fibrosis including FCGR-mediated phagocytosis and platelet homeostasis (Fig. 5E). DEGS in cluster 5, underrepresented in mWSD-exposed livers, included the predicted inactivation of senescence, natural killer (NK) cell signaling, PPAR, HIPPO, and CD40 pathways (Fig. 5F; Supplementary Table 14), consistent with decreased endothelial activation and antigen presentation^48,49,50^. Genes in these clusters predicted the activation of pathways related to protein translation (EIF2, translation initiation, termination), pathways promoting autophagy (aggrephagy, mitophagy, pexophagy, UFMylation signaling), and pathways promoting resolution of oxidative stress including cytoprotection by HMOX1 and KEAP1/NFE2L2 (Fig. 5F). Overall, RNA profiles in endothelial cells from mWSD-exposed livers predict LSEC expansion and immune recruitment and tolerance, with reduced vascular endothelial function.

**Figure 5.**
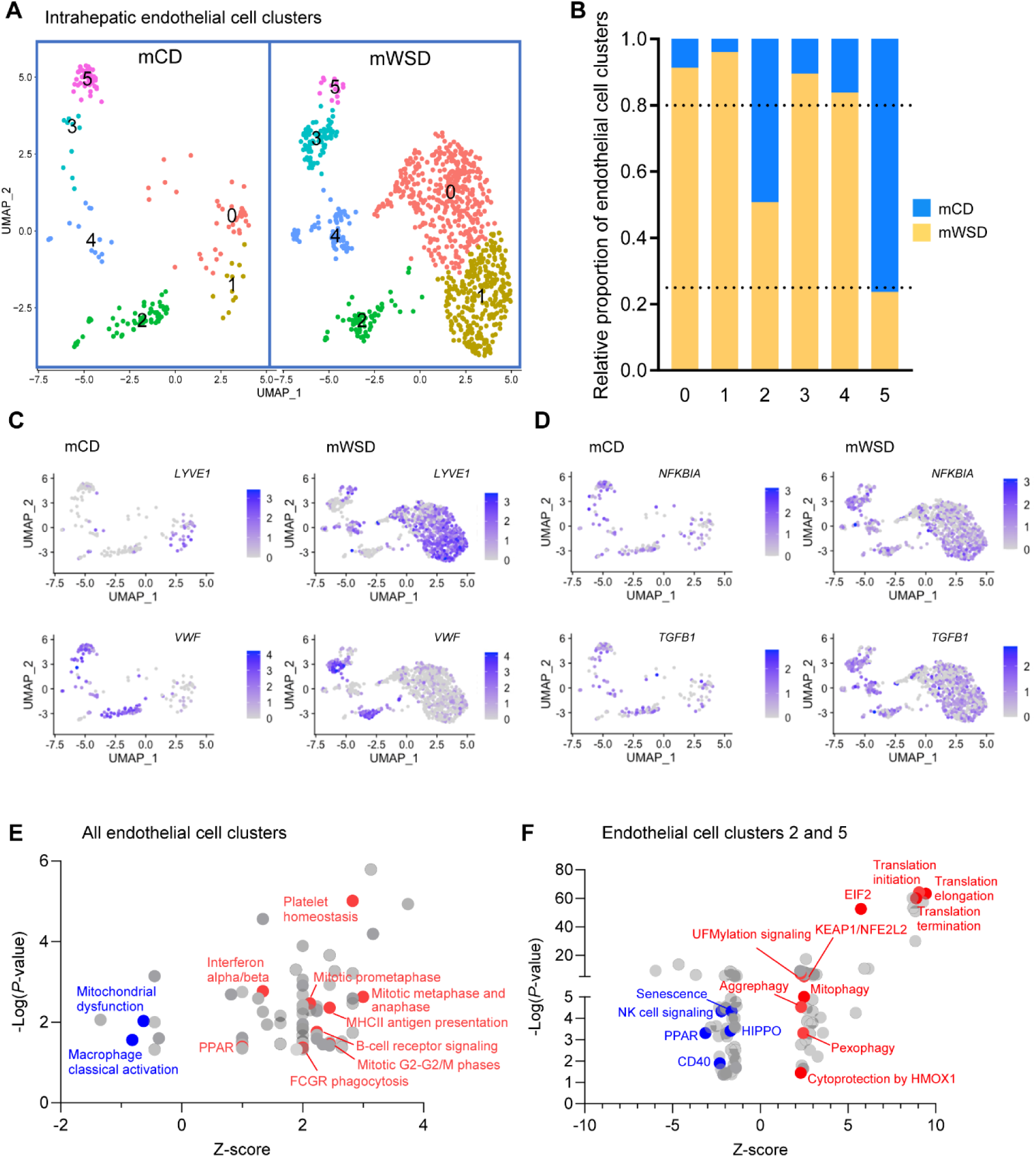
mWSD exposure is associated with preferential expansion of LSECs. **A** UMAP of endothelial cells split by maternal diet showing clusters numbered 0-5. **B** Relative proportion of each endothelial cell cluster in mCD and mWSD groups. Dashed lines represent boundaries to show decreased (lower line) and increased (upper line) proportions in mWSD- vs. mCD-exposed endothelial cells. **C** UMAP of total endothelial cells showing distribution of expression of *LYVE1*, a marker of liver sinusoidal endothelial cells, and *VWF*, a marker of vascular endothelium. **D** UMAP of total endothelial cells showing distribution of expression of *NFKBIA and TGFB1* by maternal diet. **E** Activation Z-scores for all the significant inactivated (7) and all activated (63) pathways (by *P*-value) derived from comparison of mWSD- vs. mCD-exposed total endothelial cells. **F** Activation Z-scores for the significant top 50 inactivated and top 50 activated pathways (by *P*-value) derived from genes defining endothelial cell clusters underrepresented in mWSD-exposed livers (clusters 2 and 5). **C**, **D** Light purple corresponds to lower expression; dark purple, higher expression. **E**, **F** Select red pathways indicate positive Z-score and activation; blue pathways, negative Z-score and inactivation; grey, top activated or inactivated pathways not highlighted.

### mWSD-exposed liver mononuclear cell metabolite profiles support impaired fatty acid oxidation

To further determine whether mWSD exposure remodeled metabolism across immune cell populations, we performed metabolomic analysis in isolated liver mononuclear cells from a subset of 3-year-old offspring from our NHP model (Supplementary Table 15). Upregulated metabolites were associated with pentose phosphate pathway, one-carbon and glutathione metabolism, nucleotides, and glycolysis, while several amino acids were downregulated (Fig. 6A). Pathway analysis supported effects on increased pentose phosphate pathway and reduced amino acid biosynthesis in mWSD-exposed immune cells (Fig. 6B). Interestingly, several lipid metabolites were decreased. Thus, increased glycolytic and pentose phosphate metabolism may compensate for reduced ATP production by free fatty acid oxidation and oxidative phosphorylation. In support, among oxidative phosphorylation genes differentially expressed by maternal diet per cell type, B cells from mWSD-exposed livers had 7/8 genes downregulated, T cells had 3/6 downregulated, macrophages had 4/6 downregulated, KCs had 7/8 downregulated, and DCs had 1/1 downregulated (Fig. 6C), suggesting reduced oxidative phosphorylation across mWSD-exposed liver immune cells.

**Figure 6.**
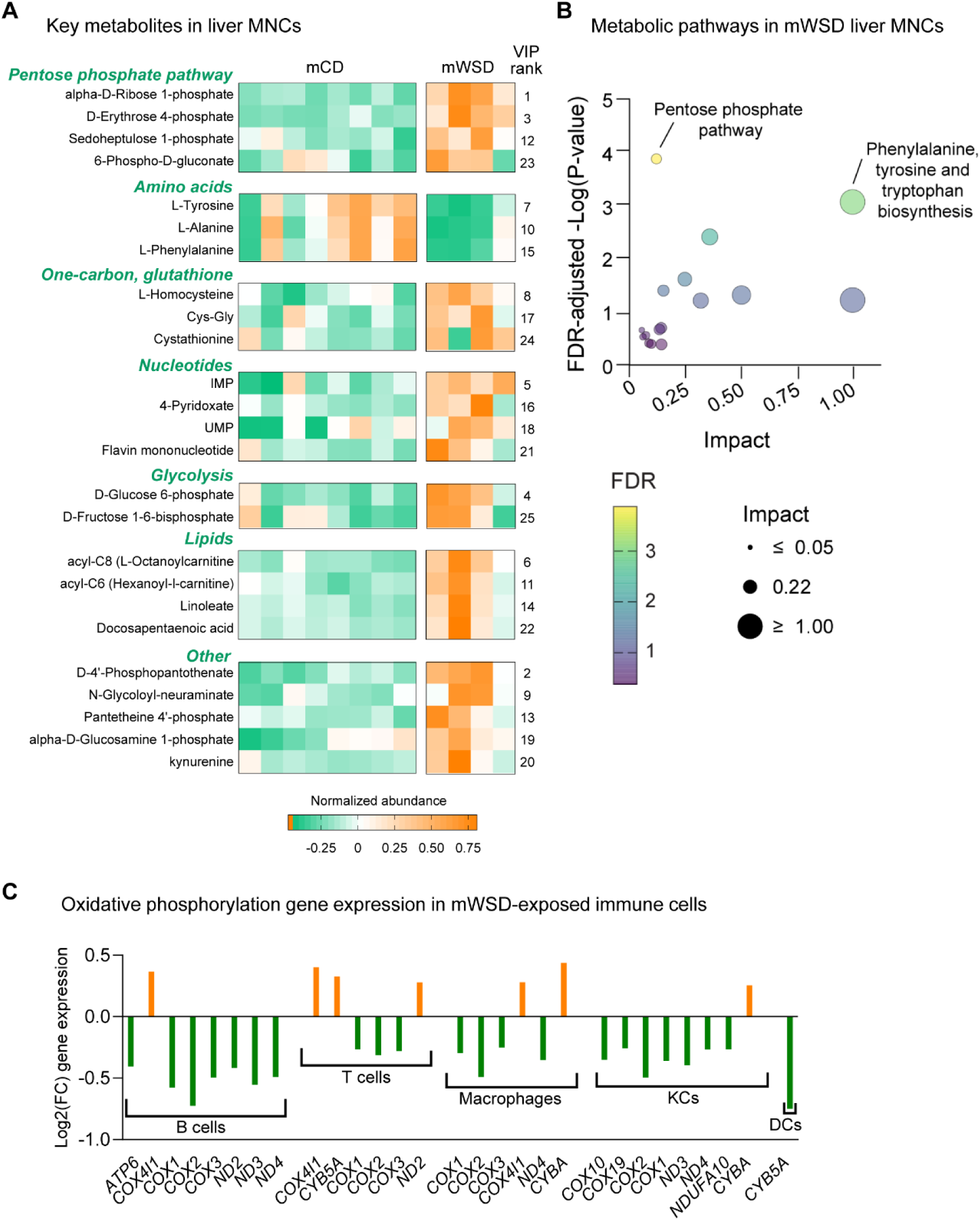
mWSD-exposed liver macrophages lack an augmented inflammatory response and liver mononuclear cells have metabolic profiles consistent with impaired fatty acid oxidation. Semi-targeted metabolomics was performed in mCD- and mWSD-exposed liver mononuclear cells (MNCs). n = 8 mCD, n = 4 mWSD. **A** Heatmap of the top 25 metabolites with the highest VIP scores from PLS-DA, with VIP ranks shown, and (**B**) enriched metabolic pathways from these metabolites, excluding the four lipid metabolites. FDR-adjusted –Log(*P*-value) and impact scores are shown for top 25 pathways. Pathways with FDR-adjusted *P* <0.05 are indicated. **C** Expression of oxidative phosphorylation genes in each population of mWSD- exposed immune cells from the scRNA-seq dataset. Orange, upregulated compared with mCD; green, downregulated. Genes shown are significantly different between mCD and mWSD groups in each cell type. n = 3 mCD, n = 3 mWSD.

### LSECs closely associate with immune cells in mWSD-exposed liver periportal regions

As LSECs are integral to the liver’s immune functions, particularly in the periportal areas^51^, we assessed the localization and abundance of different cell populations from a subset of 3-year-old offspring from our NHP model (Supplementary Table 16; Supplementary Fig. 5). CD45 immunofluorescence was used to identify all immune cells. RNAscope was used to identify additional cell types with probes targeted to *MARCO*, a marker of portal-associated fibrogenic macrophages^32^, *TIMP1*, a marker of macrophages associated with stellate cell activation^11^, and *CD37*, a gene with myriad functions playing roles in antigen presentation and immune regulation^52^, which was upregulated in macrophages, B cells, T cells, and endothelial cells from mWSD-exposed livers, and *LYVE1*, a marker of LSECs^46^.

Across whole tissue sections of juvenile liver, we found no difference in the number of CD45^+^ immune cells between maternal diet groups (Supplementary Fig. 6A; *P* = 0.6394). A trend for a modest increase in CD45^+^ cells expressing *MARCO* was observed (Supplementary Fig. 6B; *P* = 0.1485). No differences in the populations of CD45^+^ cells expressing either *CD37* or *TIMP1* were observed (Supplementary Fig. 6C, D, respectively; *P* >0.3).

Next, we looked at the frequency of CD45^+^ immune cells neighboring *LYVE1*^+^ LSECs near portal triads and central veins (Fig. 7A). The proportion of CD45^+^ cells present increased in mWSD-exposed livers in both portal triad (Fig. 7B; *P* = 0.0282) and central vein regions (Fig. 7C; *P* = 0.0464). No differences in the proportion of *LYVE1*^+^ cells in portal triad and central vein regions were observed (Fig. 7D, E, respectively; *P* >0.5). The proportion of CD45^+^ cells interacting with *LYVE1*^+^ cells increased in mWSD-exposed livers in the portal triad (Fig. 7F; *P* = 0.0049) but not the central vein (Fig. 7G; *P* = 0.2355) regions. Thus, increased interactions in the periportal sinusoids between immune cells and *LYVE1* expression (a marker of LSECs) may underlie the periportal specificity of increased collagen deposition in mWSD-exposed offspring^10,11^.

**Figure 7.**
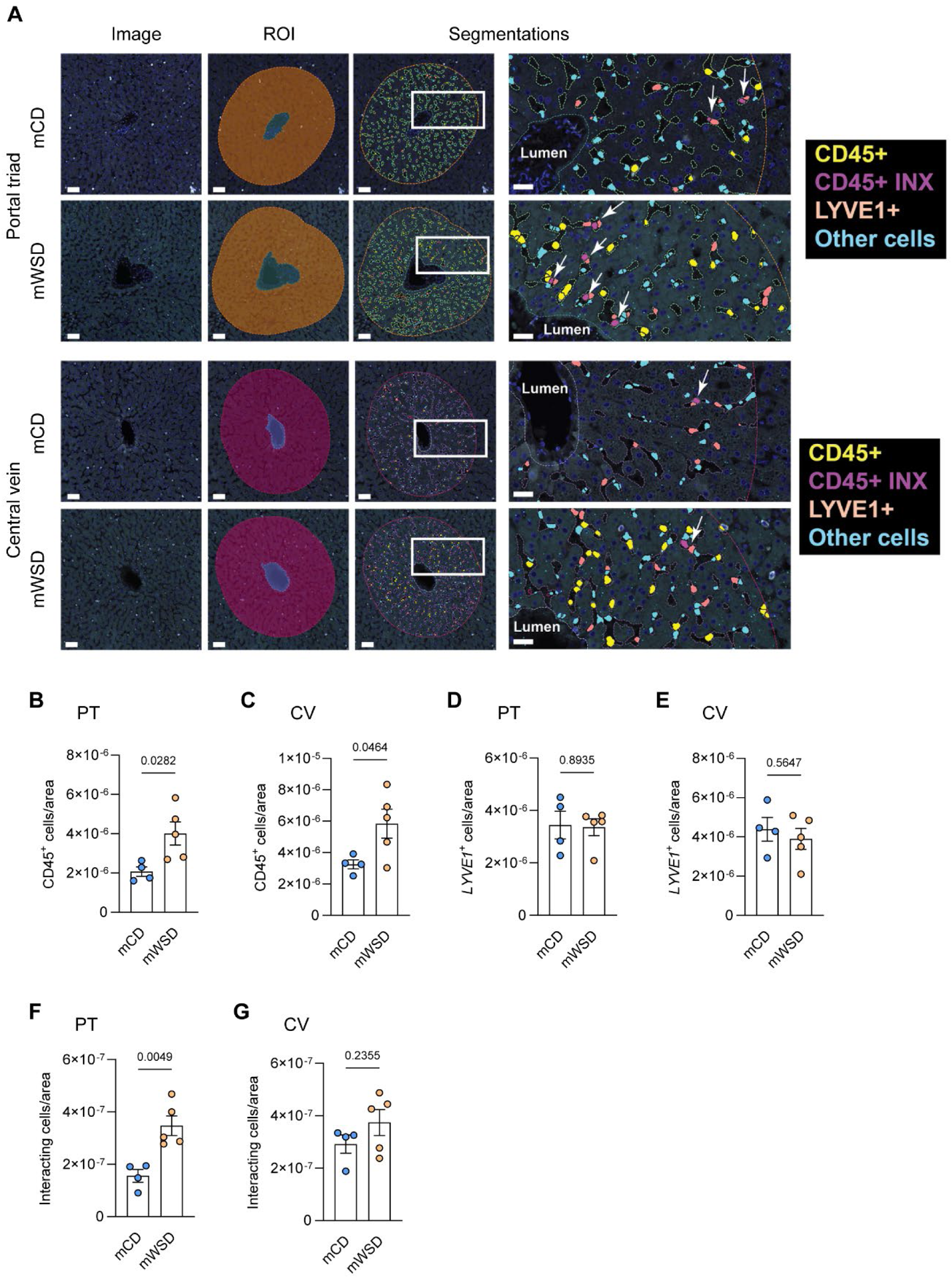
LSECs are closely associated with immune cells in mWSD-exposed liver periportal regions. **A** Representative images of portal triad (PT) and central vein (CV) regions from NHP livers depicting CD45^+^ immune cells identified by antibody (yellow), RNA probe signal for *LYVE1* via RNAscope (pale pink), CD45^+^ cells interacting with *LYVE1*^+^ cells (purple; CD45^+^ INX [interaction]), and DAPI stained cells without CD45 or *LYVE* signals cells (blue). White arrows indicate CD45^+^ cells interacting with *LYVE1*^+^ LSECs. Scale bars: 50 μm on whole PT and CV images, 20 μm on zoomed in images (right column). ROI, region of interest. Quantification of CD45^+^ immune cells as a proportion of total cells present within sinusoids in PT (**B**) and CT (**C**) regions. Quantification of *LYVE1*^+^ LSECs present within sinusoids in PT (**D**) and CT (**E**) regions. Quantification of CD45^+^ cells within 2 µM radius of an LSEC (defined as interaction between cells) within sinusoids in PT (**F**) and CT (**G**) regions. Cell counts were normalized to sinusoidal area. Each data point represents an average of 10 representative images from PT or CV regions per animal. Data are mean ± SEM. n = 4 mCD, n = 5 mWSD. *P* values are shown as analyzed by Student’s *t* test.

### Cell–cell communication analysis predicts dampened pro-inflammatory hepatic LSEC crosstalk with immune cells and extracellular remodeling in mWSD-exposed offspring

.Given the critical role of LSECs in crosstalk with other immune cells to regulate immunity and fibrotic processes, we used CellChat^53^ to predict probable cell-cell communication. This analysis predicts interactions between sender – receiver cell types by evaluating expression levels of known ligand – receptor pairs. Overall, mWSD-exposed liver displayed a dampening of crosstalk based on predicted cell-cell communication events, most notable sent from LSECs and VECs (i.e., senders) to macrophages, KCs, and T cells (i.e., receivers) compared with mCD-exposed liver (Fig. 8A). DCs displayed increased communication with KCs and B cells, while KCs showed dampened crosstalk with T cells, macrophages, and B cells.

**Figure 8.**
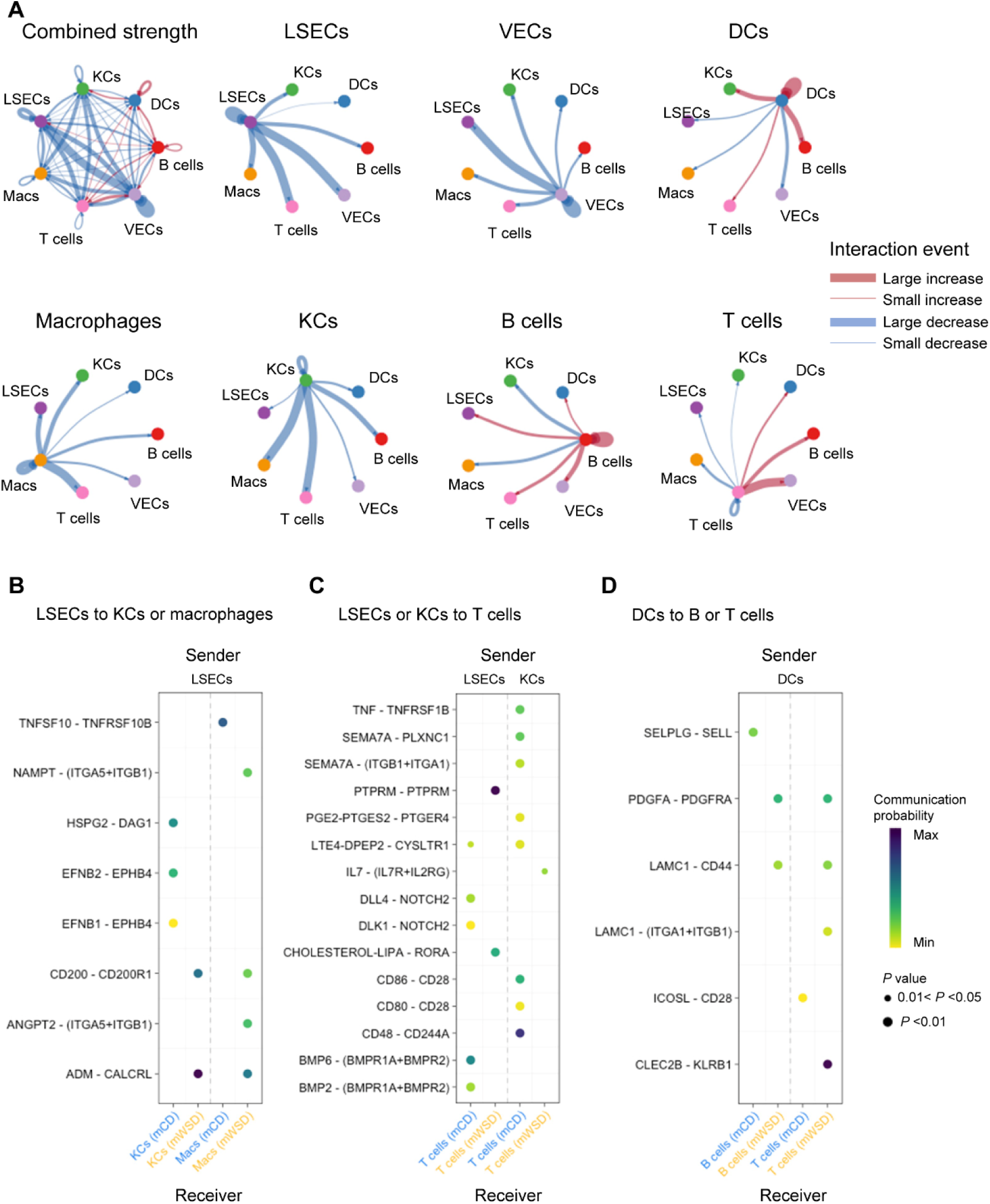
mWSD exposure drives immune dysregulated cell communication in LSECs. **A** Circle plots showing differential strength of interactions between sender and receiver cell type combinations in mWSD-exposed livers compared with mCD-exposed livers. Cell types arranged on the outside with connecting lines indicating increased (red) or decreased (blue) communication strength (indicated by the width of the connecting lines; thicker lines, increased strength of connections). Ligand – receptor pairs that are present/absent in sender – receiver pairs by maternal diet in LSECs to KCs and macrophages (Macs) (**B**), in LSECs and KCs to T cells (**C**), and in DCs to B and T cells (**D**). Presence/color of dot in viridis palette is the communication probability as a scaled model score (no dot, no communication, = 0; yellow dot, minimum communication, > 0; dark blue, maximum communication, ∼1); size of dot is the *P* value (all are *P* < 0.05 by Wilcoxon test).

Interestingly, despite decreases in overall crosstalk, LSECs, as the sender, showed increased predicted signaling to KCs and macrophages, as receivers, in mWSD-exposed livers specifically through the CD200 – CD200R1 ligand – receptor pair, involved in suppressing inflammation and phagocytosis^54^, and the ADM (adrenomedullin) – CALCRL (calcitonin receptor-like receptor) pair (Fig. 8B; Supplementary Fig. 7A). LSECs also showed increased predicted signaling to macrophages in mWSD-exposed livers through ITGA5+ITGB1 (α5 and β1 integrins), cell surface receptors responsible for adhesion to endothelial cells^55^ (Fig. 8B). Conversely, in LSEC to KCs, HSPG2 – DAG1, a pair that is responsible for ECM maintenance^56^, and EFNB1/2 (ligands ephrin B1 and B2) – EPHB4 (their cognate receptor), which modulates pathways for cell adhesion and migration^57^, were increased in mCD-exposed livers compared with mWSD-exposed livers (Fig. 8B). TNFSF10 – TNFRSF10B crosstalk in LSECs to macrophages was increased in mCD-exposed livers compared with mWSD-exposed livers (Fig. 8B). Thus, the overall message sent from LSECs to KCs and macrophages in the mWSD-exposed liver is one of vascular destabilization, immunosuppression, and pro-fibrotic signaling.

We also examined the communication patterns from LSECs or KCs as senders to T cells as the receiver. CD48, CD86, SEMA7, TNF, NOTCH, and CD80 crosstalk were present in mCD-exposed livers, among others, but absent in mWSD-exposed offspring (Supplementary Fig. 7B). In mWSD-exposed livers with LSECs as the sender, PTPRM – PTPRM signaling, which promotes cell adhesion^58^, and LIPA – RORA crosstalk were increased compared with mCD-exposed livers and DLL4 – NOTCH2 was increased in mCD-exposed livers and absent in mWSD-exposed livers (Fig. 8C). Among the ligand – receptor pairs that were increased in mCD-exposed livers and absent in mWSD-exposed livers with KCs as the sender, CD86 – CD28 and CD80 – CD28 have been shown to influence the homeostasis of Tregs^59^ (Fig. 8C). Further, CD48 – CD244A, SEMA7A – ITGB1/A1, and TNF – TNFRSF1B, regulators of the inflammatory responses of lymphocytes^60,61,62^, were undetected in mWSD-exposed livers compared with mCD-exposed livers (Fig. 8C). Taken together, these parings reveal distinct ligand – receptor signaling patterns reflecting shifts in immune responses and point to an inability to mount an acute adaptive immune response in mWSD-exposed livers.

Activated DCs in MASLD can lead to a dysregulated T cell response. Communication from DCs as the sender to B or T cells as receivers was increased in mWSD-exposed liver. LAMININ, PDGF, and CLEC pathways were present in mWSD-exposed livers and SELPLG and ICOS were absent (Supplementary Fig. 7C). The ligand – receptor combinations PDGFA – PDGFRA, important for IL-10 production, and LAMC1 (laminin subunit gamma 1) – CD44 were increased in B and T cells from mWSD-exposed livers (Fig. 8D). LAMC1 – ITGA1/B1 and CLECB2 – KLRB1 (C-type lectin-like receptor 2-CD161) were increased in mWSD-exposed liver (Fig. 8D). SELPLG (selectin P glycoprotein ligand) – SELL (L-selectin), involved in immune cell trafficking during inflammation^63^, was increased in B cells and ICOSL (inducible T cell costimulator ligand) – CD28, involved in T cell activation^64^, was increased in T cells from mCD-exposed livers compared with mWSD-exposed livers (Fig. 8D). Thus, communication between DCs and B and T cells favored IL-10 production and tolerance, outweighing activating inflammatory signals.

## Discussion

Our prior studies in NHP fetal and juvenile offspring exposed to mWSD^10,11,29,30^, together with observations in pediatric MASLD patients^6^, reveal a consistent predominance of hepatic periportal fibrosis and portal immune cell recruitment^65^, rather than the classic perisinusoidal pattern seen in adults. Our current results suggest that maladaptive immune responses and metabolic reprogramming evolve from early mWSD exposure leading to increased immune tolerance in juvenile offspring. Importantly, such changes appear before more advanced fibrosis occurs, indicating that immune signaling alterations are closely tied to early MASH progression in our model of pediatric MASLD.

We identified RNA and metabolic signatures supporting innate and adaptive immune cell tolerance rather than inflammation, highlighted by disruption of the normal cell-cell communication that controls inflammation and tissue architecture (Fig. 9). This demonstrates that mWSD exposure promotes pathological fibrosis early in development (i.e., perinatal from in utero through lactation), despite switching to a normal chow diet at weaning. Reduced immune-immune cell crosstalk or missing costimulatory signals lead to immune tolerance. Tolerogenic DCs, increased in mWSD-exposed livers herein, can induce the differentiation of T helper cells into FOXP3-expressing Tregs, which promote tolerance. We previously reported expansion of DCs in mWSD-exposed fetal livers with a reduction in transitional effector memory T cells^11^, suggesting that impaired DC-mediated immune training and/or tolerance begins in utero. Since lymphoid-derived DCs are capable of T cell activation, early and persistent intrahepatic DC tolerance and block of maturation by mWSD exposure may be key factors reducing the adaptive immune system’s response to oxidative stress, allowing for oxidative stress and fibrosis to occur unchecked (i.e., unresolved) in our model of mWSD exposure-associated MASLD.

**Figure 9.**
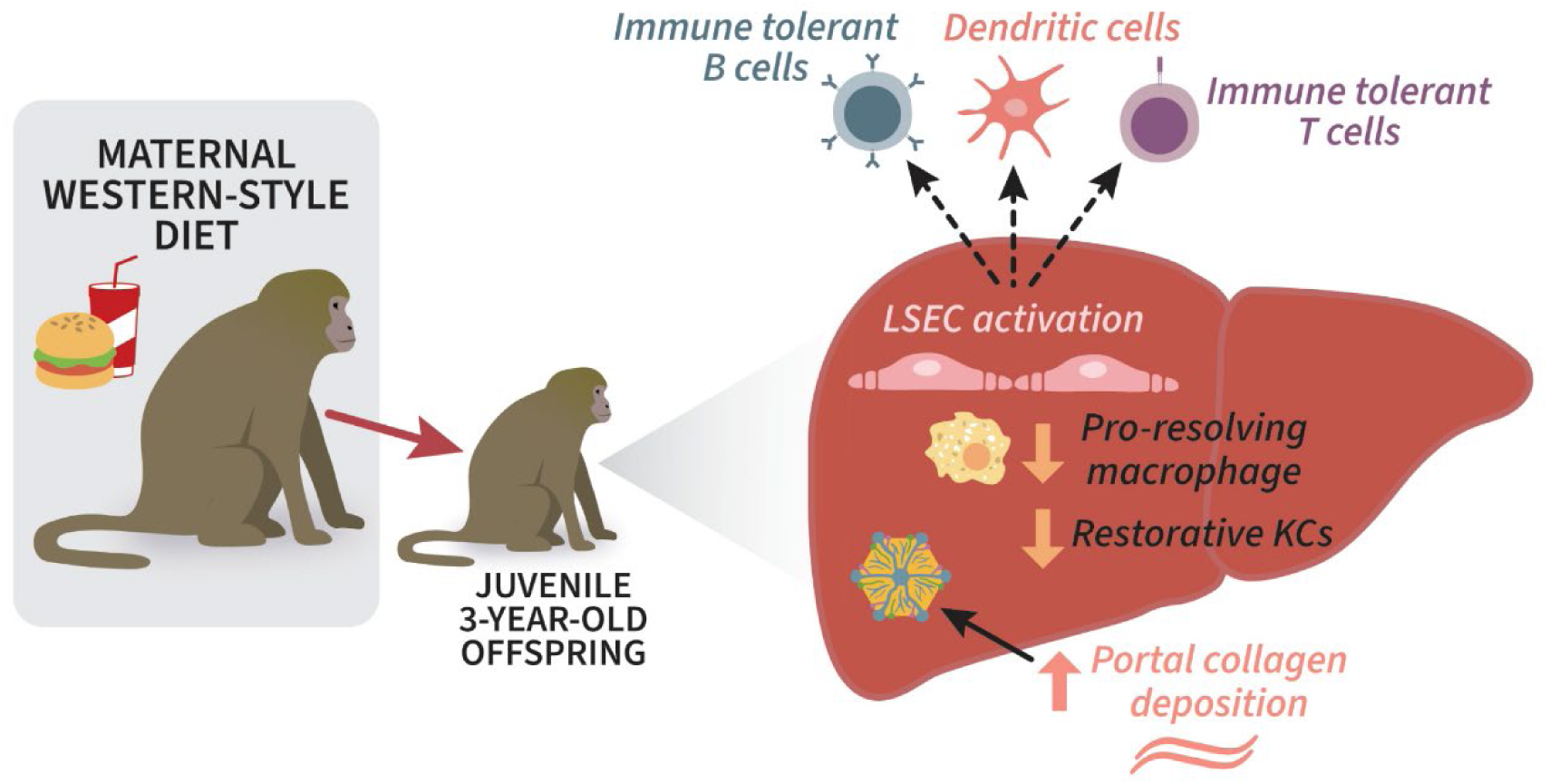
mWSD promotes immune tolerance and LSEC dysfunction in NHP juvenile offspring liver. Exposure to an mWSD revealed altered hepatic liver sinusoidal endothelial cells (LSECs) and innate and adaptive immune landscapes in juvenile offspring, specifically affecting macrophages, B cells, dendritic cells, and T cells that together promote immune tolerance. We observed a marked reduction in immune-to-immune cell crosstalk, driven primarily by a lack of essential costimulatory signals between LSECs and both innate and adaptive immune cell populations. While Kupffer cells (KCs) were diminished, infiltrating macrophages adopted a non-inflammatory, non-reparative phenotype characterized by the upregulation of genes involved in stellate cell interactions. Expanded and activated LSEC-immune cell interactions and LSEC-associated increases allow for ongoing immune tolerance and oxidative stress but are permissive of fibrogenesis in the portal zone without overt inflammation. Expanded and activated LSECs sustain a local tolerogenic microenvironment; this immune-suppressive state is permissive of portal zone fibrogenesis that resists postnatal dietary correction.

T cell-mediated inflammation links steatosis and steatohepatitis, in part by triggering B cell-mediated antibody production and macrophage infiltration, supporting a role for T cells as a central driver in MASLD/MASH pathogenesis^66^. Our data showed reduced T cell effector function and evidence of a naïve T cell phenotype in mWSD-exposed livers. Particularly, mWSD-exposed T cells showed decreased inflammation (IL-6, NFκB, IL-8), increased apoptotic signaling (death receptor, apoptosis), and activation of translation machinery without transcriptional activation. Although DCs were relatively depleted in mWSD-exposed livers, examining DCs as the sender with B or T cells as receivers, overall communication was increased in mWSD-exposed liver. Using CellChat, we found that the LAMC1 – ITGA1/B1 ligand – receptor pair, important for DC-T cell interactions and potentially ECM remodeling^67^ was increased in mWSD liver, while ICOSL – CD28, important for T cell activation^64^, was absent in mWSD-exposed T cells compared with mCD-exposed T cells. PDGFA – PDGFRA, important for IL-10 production, and CLECB2 – KLRB1, important for T cell function^68^ were increased in mWSD-exposed T cells. The interaction between SELPLG and SELL, which is responsible for immune cell trafficking through initial adhesion of leukocytes to endothelial cells during inflammation^63^, was absent in mWSD-exposed B cells.

Among the significantly differentially regulated ligand – receptor pairs important for Treg activation, with KCs as the sender, CD86 – CD28, CD80 – CD28, CD48 – CD244A, and SEMA7A – (ITGB1/A1) were lacking in mWSD-exposed T cells. Further, TNF – TNFRSF1B from KCs and DLL4 – NOTCH from LSECs, which both regulate the inflammatory responses of T lymphocytes^62,69^ were also absent in mWSD-exposed T cells. LAMININ, PDGF, and CLEC2B pathways, critical for trafficking T and B cell movement and acting as inhibitors of T cell function^70^ and T cell tolerance^71^, were predicted to be increased in mWSD-exposed liver. In healthy conditions, T cells and DCs respond to acute oxidative stress by triggering acute inflammation and resolution^72,73^, but chronic exposure to oxidative stress can lead to their dysfunction and a state of anergy. Previously, we have shown increased oxidative stress in mWSD-exposed fetal livers^11^ and juvenile livers^10^, yet the tolerant phenotypes remained in the dataset herein. Together, these observations suggest a breakdown in adaptive immune regulation, indicating that livers exposed to mWSD with associated persistent oxidative stress are unable to mount a competent T cell response.

Surprisingly, a large increase in B cells in mWSD-exposed liver was observed, with a predicted inactivation of TNF and IL-3 inflammatory and metabolic pathways. B cells drive liver fibrosis in MASH^74,75^ and inaction of some B cell subsets is associated with MASH severity^66^. Importantly, mWSD exposure was associated with the emergence of a unique B cell cluster (cluster 4), in which IL-10 and VEGF were increased, suggesting an influence on anti-inflammatory and local vascular remodeling functions. Selective expansion of novel immunosuppressive B cell clusters, possibly representing Bregs, and polarization of B cells towards tolerogenic phenotypes may be a unique feature of our model of mWSD exposure. B cells that fail to properly transfer antigen or provide signals to macrophages/innate antigen presenting cells and macrophage activation/polarization may skew toward a dysfunctional or pro-fibrotic state rather than one supporting healthy tissue repair or immune surveillance.

Macrophages from mWSD-exposed livers were characterized by upregulation of stellate cell interaction genes including *MSRB1* and *TIMP1*, and a non-inflammatory, non-reparative phenotype. The increased expression of *MSRB1* and *TIMP1* in clusters of mWSD-exposed macrophages, and activation of the pro-fibrotic TWEAK pathway, supports increased interactions between macrophages and stellate cells^76,77^. Notably, TWEAK pathway activation amplifies inflammation, promotes tissue damage, and potentially impedes endogenous repair mechanisms^78^. Importantly, we did not see the emergence of a pro-fibrotic macrophage cluster unique to mWSD-exposed livers, consistent with data in adult human MASH^32^. KC loss in MASH and replacement by infiltrating monocytes promotes liver damage, fibrosis, and other features of MASH^79^. Circulating monocytes can differentiate into M2-like reparative macrophages, facilitating the resolution of inflammation and tissue repair^17,80^. However, the remaining mWSD-exposed KCs had a similar phenotype as other mWSD-exposed intrahepatic macrophages, supporting an early pro-fibrotic MASH phenotype due to loss of “restorative” KCs by mWSD exposure. mWSD-exposed macrophages had activation of complement, associated with MASLD progression^81^, and activation of death receptor and type 1 interferons^82^, consistent with the lack of reparative function. mWSD-exposed liver macrophages also had predicted inactivation of classical macrophage activation, including IL-1, IL-6, and phagocytosis, suggesting a lack of acute inflammatory response, a key to eventual tissue healing and regeneration^16^. These observations suggest that mWSD exposure remodels liver macrophages to a pro-fibrotic yet blunted pro-inflammatory phenotype, allowing for continuous liver damage and collagen deposition.

We found an increased number of physically associated LSECs and immune cells in portal regions identified by IHC and RNAscope in mWSD-exposed livers. LSECs promote immune tolerance across both the innate and adaptive immune system^83^. In MASH, KCs and recruited monocytes are driven toward an inflammatory or scar-associated phenotype based on macrophage niche occupancy and LSEC interactions^18^. In our dataset, mWSD exposure remodeled transcriptional phenotypes across all macrophage clusters studied. An intriguing hypothesis is that the expansion of LSECs disrupts liver microenvironmental niches, leading to a tolerant phenotype in macrophages despite ongoing oxidative stress and collagen deposition^10^. In support, the EFNB1 receptor signaling pathway is required for endothelial tight junction integrity^84^ and was absent in LSEC to KC communications from mWSD-exposed offspring. On the other hand, CD200, a cell surface glycoprotein that functions as an immunosuppressive and anti-inflammatory molecule by interacting with its receptor, CD200R^85^, was increased in LSECs to KCs and macrophages crosstalk in mWSD-exposed livers. Expression of ADM and its receptor, CALCRL, was also increased between LSECs to KCs and macrophages in mWSD-exposed liver and is involved in immune tolerance and dampening inflammation^86^. The expansion of LSEC subsets is a key early factor in MASH initiation and strongly associated with MASH severity^87^. In adult rodent models of MASLD, LSEC dysfunction precedes fibrosis^88,89,90^, while inhibiting LSEC maladaptation via eNOS activators in methionine- and choline-deficient diet-induced MASH mouse models^91^ or inhibiting epigenetic reprogramming in a minipig MASH model^92^ limited fibrogenesis, indicating an early role for LSECs in triggering and maintaining MASH. Whether LSEC disruption is initiated in utero and advanced by mWSD exposure during lactation (e.g., due to microbiome influences) is unknown.

Overall, our data demonstrates that mWSD-exposed LSEC-immune cell interactions were physically increased, but important costimulatory or pro-inflammatory pathways with many functions that may reduce tolerance such as TNF were missing, whereas other tolerance-promoting pathways like CD200 were increased in mWSD-exposed LSECs, thus allowing for pathological tolerance. We speculate that mWSD exposure-associated oxidative stress reduces LSECs ability to produce and handle nitric oxide and allows for increased but incomplete (lacking important costimulatory/pro-inflammatory pathways) LSEC-immune cell interactions and LSEC-associated increases in B cells/antigen presenting cells^93^, thus allowing for ongoing immune tolerance but being permissive of fibrogenesis without overt inflammation, as described previously in our model^10,11,30^. Importantly, healthy LSECs prevent HSC activation and promote reversion of HSCs to quiescence, but capillarized or otherwise damaged LSECs do not^94^. Our previous work showed that mWSD-exposed livers from 3-year-old NHPs have ongoing HSC activation and fibrogenesis^30^, suggesting that the ability of LSECs to revert HSCs to a quiescent phenotype is impaired by mWSD exposure.

In our model, mWSD exposure is accompanied by reactive oxygen species production in 3-year-old NHP livers at the level of the mitochondria with ongoing oxidative stress^10^. Here, we observed downregulated oxidative phosphorylation, increased glycolysis, and impaired free fatty acid metabolism across most immune cell types, consistent with a non-reparative and non-proliferative immune phenotype^95^. In adaptive immune cells, oxidative phosphorylation is associated with cellular proliferation and effector response^96^, and in macrophages, is associated with a reparative anti-inflammatory phenotype, along with fatty acid oxidation^97^. The functional metabolic reprogramming of the immune system toward impaired oxidative phosphorylation may occur early in development in hematopoietic stem and progenitor cells, which produce all progeny immune cells, as we reported in mWSD-exposed fetal and 3-year-old NHP previously^29^.

Very little is known about the developmental pathophysiology and unique disease progression trajectories in children with MASLD^3^. Persistent LSEC dysfunction over time may set the stage for accelerated fibrosis, with transcriptional signatures and histological evidence of increased periportal immune-LSEC interactions, suggesting pathological immune tolerance to ongoing damaging stimuli in the liver (oxidative stress) and lack of repair with ongoing fibrogenesis despite consuming a healthy post-weaning diet for 2.5 years. Understanding the balance between tolerogenic and immunogenic immune states is crucial, as pediatric MASLD can progress more rapidly compared with adults, often leading to cirrhosis and transplantation in early adulthood^98,99^. A detailed study defining the early molecular changes during different stages of pediatric MASLD awaits further study. Given the severity of pediatric MASLD/MASH progression in young pre-adolescents^100^, even in the absence of obesity^1^, our results underscore the need for early detection/intervention from maternal dietary exposures driving the development of MASLD in the pediatric population.

## Methods

### Maternal WSD-induced obesity model

All animal procedures were conducted in accordance with the guidelines of the Institutional Animal Care and Use Committee of the Oregon National Primate Research Center and Oregon Health & Science University. The Oregon National Primate Research Center abides by the Animal Welfare Act and Regulations enforced by the United States Department of Agriculture. The 3-year-old juvenile offspring studies were conducted as previously reported^29^. The offspring were housed with dams on their respective diets through lactation until weaning at ∼7 months old, then maintained on CD until necropsy at 3 years of age. Dams^101^ and offspring^10^ were phenotyped as previously described.

### Liver mononuclear cell isolation

Liver mononuclear cells were obtained via perfusion and collagenase digestion of a portion of the right lobe of the liver from 3-year-old mCD- and mWSD-exposed offspring as reported previously, and livers were flushed with HBSS to remove circulating immune cells^29,31,33^. Hepatocytes were removed from the total mixture of digested cells by low-speed centrifugation at 100 x *g* for 5 min. The supernatant containing all nonparenchymal cells was filtered and spun at 800 x *g* for 10 min at 4°C to pellet nonparenchymal cells. Cells were resuspended in 24% Histodenz (MilliporeSigma), and gradients were prepared and spun at 1,500 x *g* for 20 min at 4°C with brake turned off. Cells at the interface (referred to as liver mononuclear cells) were collected and washed, and aliquots were either cryopreserved in 90% FBS/10% DMSO, stored in liquid nitrogen, and used for scRNA-seq analysis, or frozen immediately as a pellet at −80°C for metabolomics analysis.

### scRNA-seq

Single-cell RNA-seq data was analyzed as previously described^30^. Liver MNCs were stained with propidium iodide and sorted on a FACSAria Fusion flow cytometer (BD Biosciences). Pooled single-cell libraries for the mCD- and mWSD-exposed liver (n = 3 per pool per group; n = 1 female in mCD and the rest are males) were generated using the 10x Genomics Chromium Next GEM Single Cell 3’ Reagent Kit v3.1. A total of 7010 and 7399 cells were captured for the mCD and mWSD pools, respectively using the 10x Chromium Controller. Sequencing depth per sample averaged 119.5K reads per cell. Read mapping and expression analysis were conducted using the 10x CellRanger pipeline with custom Seurat scripts, and reads were aligned to the macaque genome (Mmul10). UMAPs were generated using normalized gene expression. Differential gene expression within and between clusters was analyzed using Student’s *t* tests with custom Seurat scripts. To demultiplex the two CellRanger possorted.BAM files from the original pools, cell barcodes for each of the cells remaining after QC were used to isolate RNA reads into single cell BAMs. Single nucleotide polymorphism (SNP) genotypes were generated from the single cell BAMs using FreeBayes^102^. An assignment test using the SNP genotypes for each cell was accomplished using ADMIXTURE^103^. Cell assignments to 1 of 3 animals per pool were imported to Seurat and used to identify the pooled single cells back to their animal of origin. The demultiplexed identifiers mCD1-3 and mWSD1-3 were then used throughout the rest of the analyses. Cluster markers used to delineate cell type identity are: B cells (CD19^+^), cholangiocytes (ALB^+^, F2^+^, FGA^+^, FGG^+^, SOX9^+^), DCs (CLEC4C^+^, IRF7/8^+^), endothelial cells (CDH5^+^), hepatocytes (ALB^+^, F2^+^, FGA^+^, FGG^+^, SOX9^-^), KCs (CLEC9A^+^), macrophages (CLEC4A^+^), stellate cells (COL1A1^+^), and T cells (CD2^+^). To calculate the relative proportion of clusters within each cell cluster, the abundance of cells in each cluster per maternal diet group was divided by the total cells in both diet groups.

Cell-cell communication analysis was conducted using the R package CellChat version 2.2^53^. Metadata from Seurat was imported to create a CellChat object for each maternal diet group (mCD and mWSD). We used the annotated cell populations from Fig.1, excluding hepatocytes, cholangiocytes, and stellate cells, and divided liver endothelial cells into LSECs and VECs based on their expression of *LYVE1* and *PROX1*, respectively. The human ligand-receptor database was used^104,105^; no database for NHP currently exists. CellChat infers intracellular communications by identifying over-expressed ligands and receptors in sender and receiver cells, respectively, and comparing combinations to the database. Biologically significant communications were calculated by the statistically robust ‘trimean’ method. Finally, objects were merged to directly compare maternal diet groups. Interaction strength based on the probability of communication was the metric used to compare differential interactions between groups.

### Ingenuity pathway analysis

Lists of DEGs and the linear fold change compared with the mCD group were used as inputs in IPA (Qiagen) to identify predicted pathways and putative upstream regulators modified by mWSD exposure. Canonical pathways that met a threshold of −Log(*P* value) of 1.3 or higher were declared significant. Positive Z-scores indicate predicted activation of the pathway, negative Z-scores indicate inactivation, Z-scores of 0 indicate neither activation nor inactivation, and pathways with no Z-score indicate that IPA was unable to calculate a Z-score. Predicted upstream regulators were declared significant with a *P* value of 1E-5 or lower and the list was filtered to include categories named in IPA as transcription regulator, ligand-dependent nuclear receptor, peptidase, group, kinase, growth factor, enzyme, transporter, and phosphatase. All entries classified as “chemical” and fusion gene/product were removed, except for LPS and N-acetylcysteine, which belonged to chemical toxicant and drug, respectively. The top pathways by magnitude of Z-score are presented in the figures.

### Targeted metabolomic analysis

Metabolomics analysis of liver mononuclear cells was performed using MetaboAnalyst software (V6.0). Peak intensities were normalized by median with range scaling and analyzed by PLS-DA. Samples with zero values were replaced with 1/5^th^ of the minimum value across samples for that feature. Normalized data from the top 25 metabolites with the highest variable importance in projection (VIP) scores are shown by heatmaps. Pathway enrichment analysis in the KEGG database was performed using the top 25 VIP metabolites, excluding the four lipid metabolites. Pathways were considered significant with FDR-adjusted *P* ≤0.05.

### Liver histology

Liver tissue samples from the left lobe were fixed in 10% zinc/formalin for 24 h followed by storage in 70% ethanol. Samples were paraffin-embedded and 5 μm thick sections were prepared on slides. Slides were deparaffinized in xylene followed by rehydration in a series of ethanol/water washes. Second harmonic generation (SHG) was performed as previously described^10,11^.

### RNAscope and IHC

Dual chromogenic RNAscope detection was used to perform in situ mRNA hybridization according the manufacturer’s protocol (Advanced Cell Diagnostics [ACD]) as described previously^11^. Briefly, the slides were deparaffinized, heat treated in ER2 buffer, incubated with protease enzyme, and the C1-channel probe (ACD) was applied and amplified with complimentary HRP-conjugated scaffolding. Opal 520 fluorescent reagent (Akoya) was then applied, followed by washing and subsequent application of C2, C3, and C4 probes (ACD) in the Opal 570, Opal 620, and Opal 690 channels (Akoya). Primary antibody specific for NHP CD45 was then applied, followed by rabbit-specific HRP polymer and Opal 780 (Akoya).

### Visiopharm analysis

#### Whole slide analysis

Juvenile monkey liver tissue sections were scanned using PhenoImager HT (Akoya Biosciences) at 20X (0.5 µm/pixel) magnification. The following channels were detected: Sample Autofluorescence (Sample AF), DAPI (DAPI), TIMP1 (Opal 520), LYVE1 (Opal 570), CD37 (Opal 620), MARCO (Opal 690), CD45 (Opal 780). Signals were unmixed and sample autofluorescence was subtracted using Inform software (Akoya). Files for the scanned tissue sections in “.unmixed.qptiff” format were imported into the Visiopharm^TM^ platform (Visiopharm). The scanned sections were analyzed through a series of custom-made Analysis Protocol Packages (APPs) using Visiopharm^TM^ software. Tissues were detected and segmented from the background using Decision Forest (90% accuracy) machine learning method at 2X magnification utilizing the autofluorescence signal as feature (4 random sections out of 9 were used to train the algorithm). Sections’ areas were eroded by 250 µm all around to remove regions associated with nonspecific signals due to edge effect. The remaining analysis areas were calculated. CD45 positive regions and their associated nuclei were classified using Deep Learning (U-Net) methods utilizing the CD45 (Opal 780) signal band, and DAPI band, respectively, as features at 20X magnification (6 random sections out of 9 were used to train the algorithms). CD45 positive cells expressing either *TIMP1*, *CD37*, or *MARCO* mRNAs were phenotyped using Visiopharm^TM^ Phenoplex module. Cells positivity for each expression was defined by whole range of intensities for CD45^+^ cells, and RNAs positivity was thresholded by average for top 5% pixels with brightest signal intensities for corresponding staining. Co-occurrence matrices were then generated for each slide. Manual inspection for quality control of automated image analyses of all slides was performed before final calculations.

#### Portal triad (PT) and central vein (CV) analysis

For PT versus CV analysis, similar sized PT and CV lumens (10 each) were manually annotated per section. Luminal regions were then automatically radially expanded by 200 µm. Next, PT’s and CV’s sinusoids areas (>30 µm^2^) were identified using K-means clustering (3 classes; unsupervised) machine learning method at 20X magnification utilizing the autofluorescence signal as feature. All luminal and sinusoidal cells were then quantified by first identifying nuclei using Deep Learning (U-Net) method utilizing DAPI band, as feature at 20X magnification, followed by 4 pixels (2 µm) dilation process to account for the cell’s cytoplasm. CD45^+^ and *LYVE1*^+^ cells were phenotyped semi-automatically. CD45 positivity was defined as cells whose lower 50% pixels to have an average CD45 signal. *LYVE1* positivity was defined as cells whose brightest 2% pixels to have an average *LYVE1* signal. Proportions of CD45^+^ and *LYVE1*^+^ cells from total cell counts were calculated for each PT and CV luminal and sinusoidal areas. In PT and CV sinusoids, CD45^+^ cells within 2 µm distance from *LYVE1*^+^ cells were categorized as “interacting” and their proportions from total CD45^+^ cells counts were calculated. Manual inspection for quality control of automated image analyses of all slides was performed before final calculations. Data were analyzed by Student’s *t* test.

## Data availability

Single-cell RNA-seq data have been deposited to GEO (accession number GSE310320) and are publicly available as of the date of publication.

## Acknowledgments

This work was supported by NIH grants R24-DK090964 (J.E.F., K.M.A., P.K.), R01-DK128416 (J.E.F.), F30-DK122672 (M.J.N.), R01-DK108910 (S.R.W.), P30-DK048520 (J.E.F., CU Anschutz Nutrition Obesity Research Center [NORC]), P51-OD011092 (OHSU for operation of the ONPRC), and the Advanced Fluorescent Microscopy Core grants at CU Anschutz (P30-NS048154, P30-DK116073). The authors thank Tyler Dean and Diana Takahashi at OHSU for assistance with animal studies and Jenny Gipson at the Institutional Research Core Facility at OUHC for assistance with the scRNA-seq.

## Author contributions

MJN, ED, SRW, and JEF conceived and designed the experiments. MJN, MM, RCJ, SRW, and JEF wrote and edited the manuscript. MJN, MM, and ED performed the experiments and analyzed data, unless noted otherwise. SIA designed Visiopharm Apps. MJN, MM, SIA, and ED performed histology analyses. BNN and KLJ analyzed scRNA-seq data. PK, KMA, CEM, MG, SRW, and JEF assisted with development of the NHP model and data interpretation. All authors approved the final version of the manuscript.

