## Supplementary Fig. for "Maternal Western-style Diet Promotes Immune Tolerance and Liver Sinusoidal Endothelial Cell Dysfunction in Nonhuman Primate Juvenile Offspring Liver"

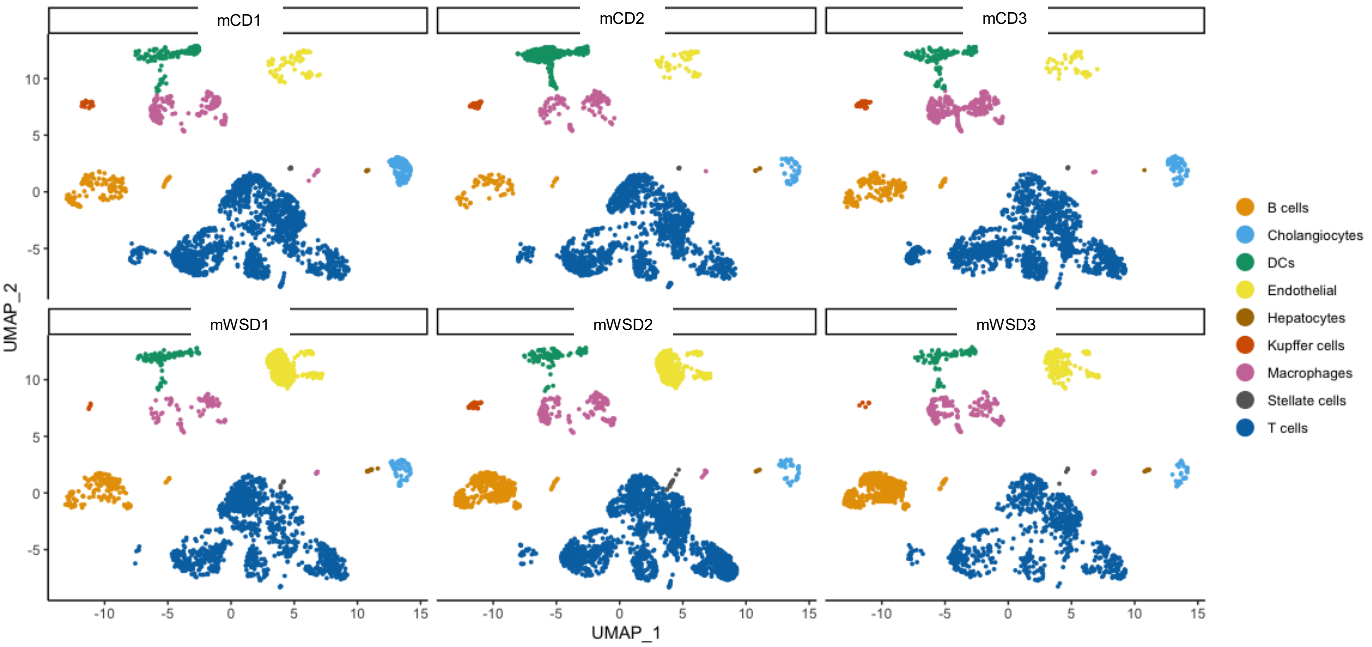


**Supplementary Fig. 1.** UMAPs of total non-parenchymal cells in liver from each individual 3-year-old NHP from scRNA-seq analysis. n = 3 offspring per group.

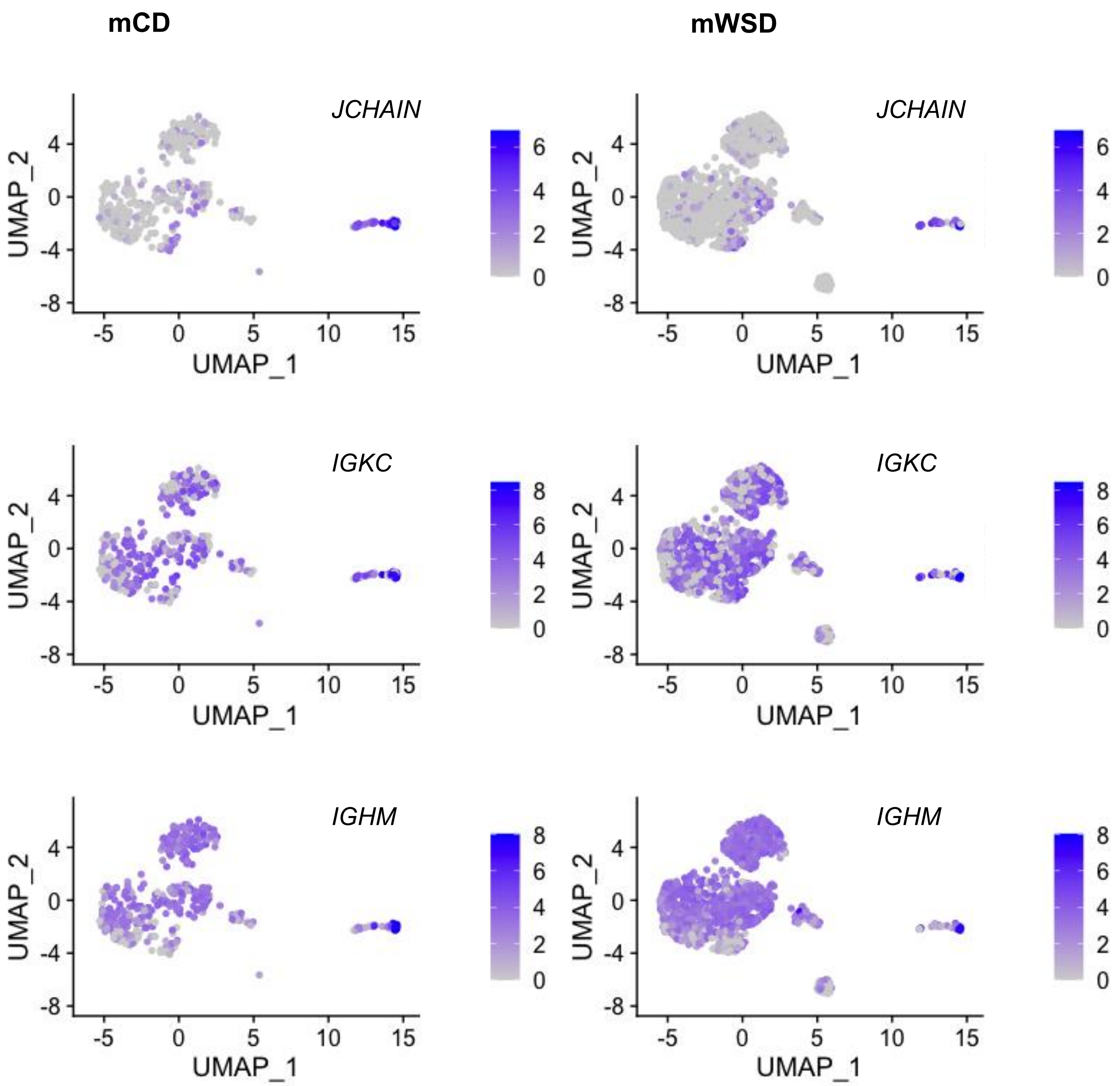


**Supplementary Fig. 2.** Markers of plasma cells in B cell clusters in mCD- and mWSD-exposed livers. Light purple corresponds to lower expression; dark purple, higher expression.

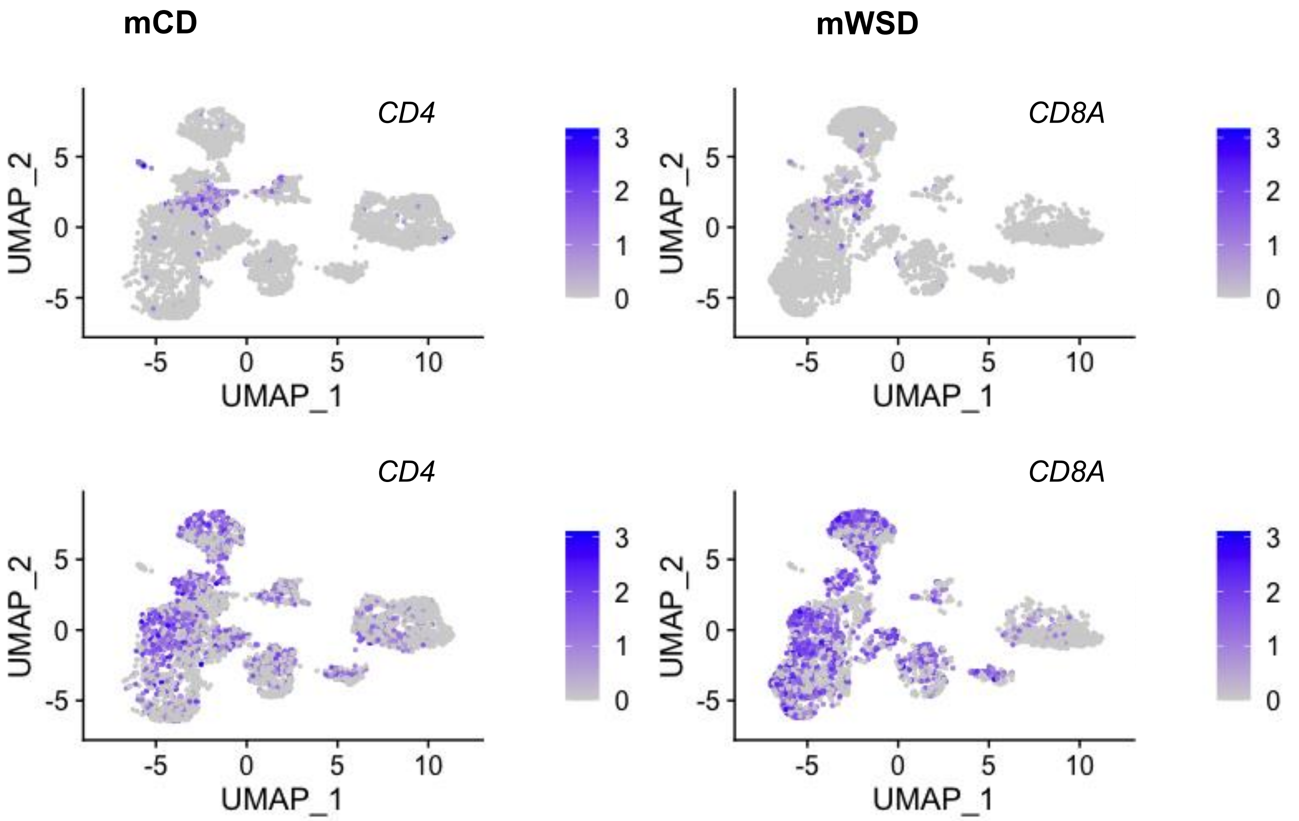


**Supplementary Fig. 3.** UMAP of CD4 and CD8 expression in T cells in mCD- and mWSD-exposed liver MNCs. Light purple corresponds to lower expression; dark purple, higher expression.

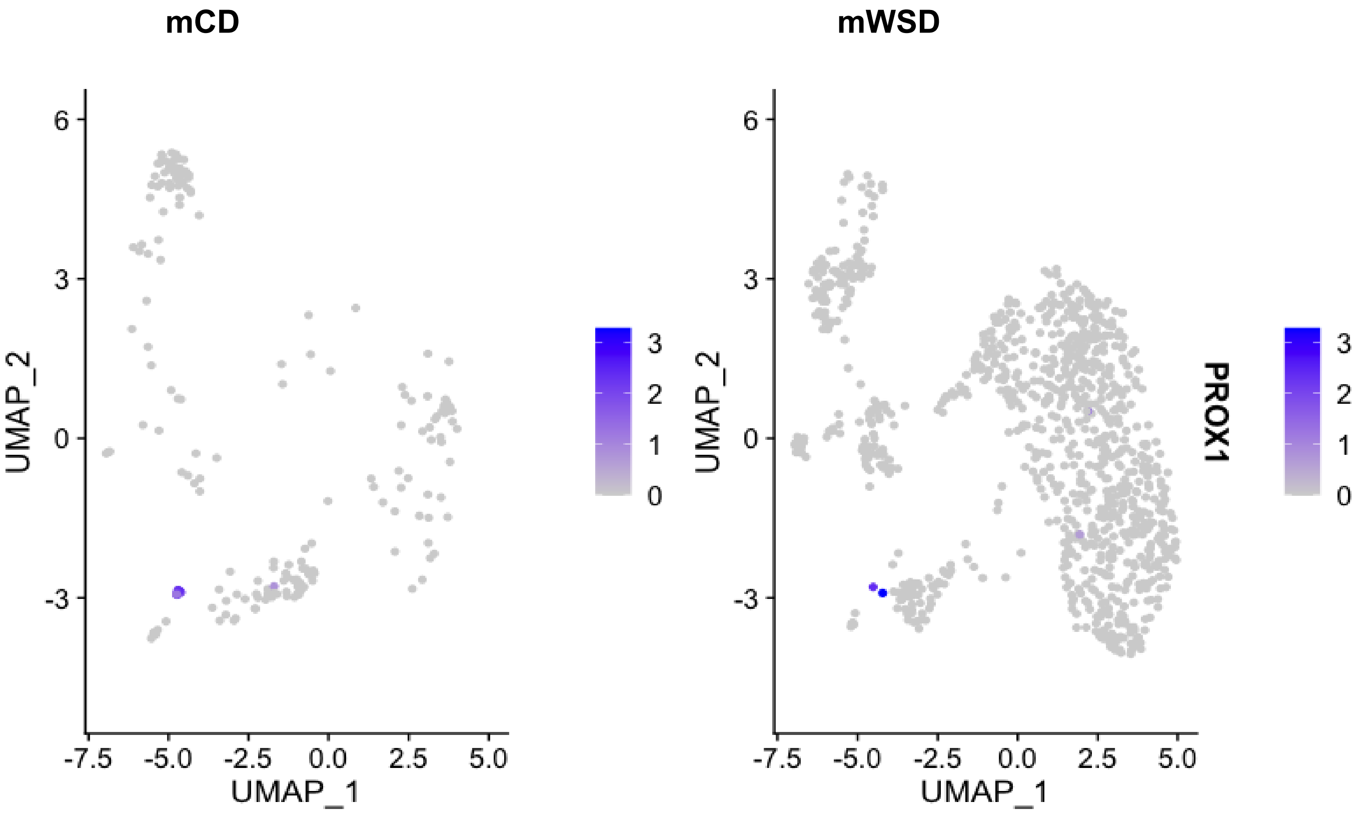


**Supplementary Fig. 4.** *PROX1* expression in intrahepatic endothelial cells in mCD- and mWSD-exposed liver MNCs. Light purple corresponds to lower expression; dark purple, higher expression.

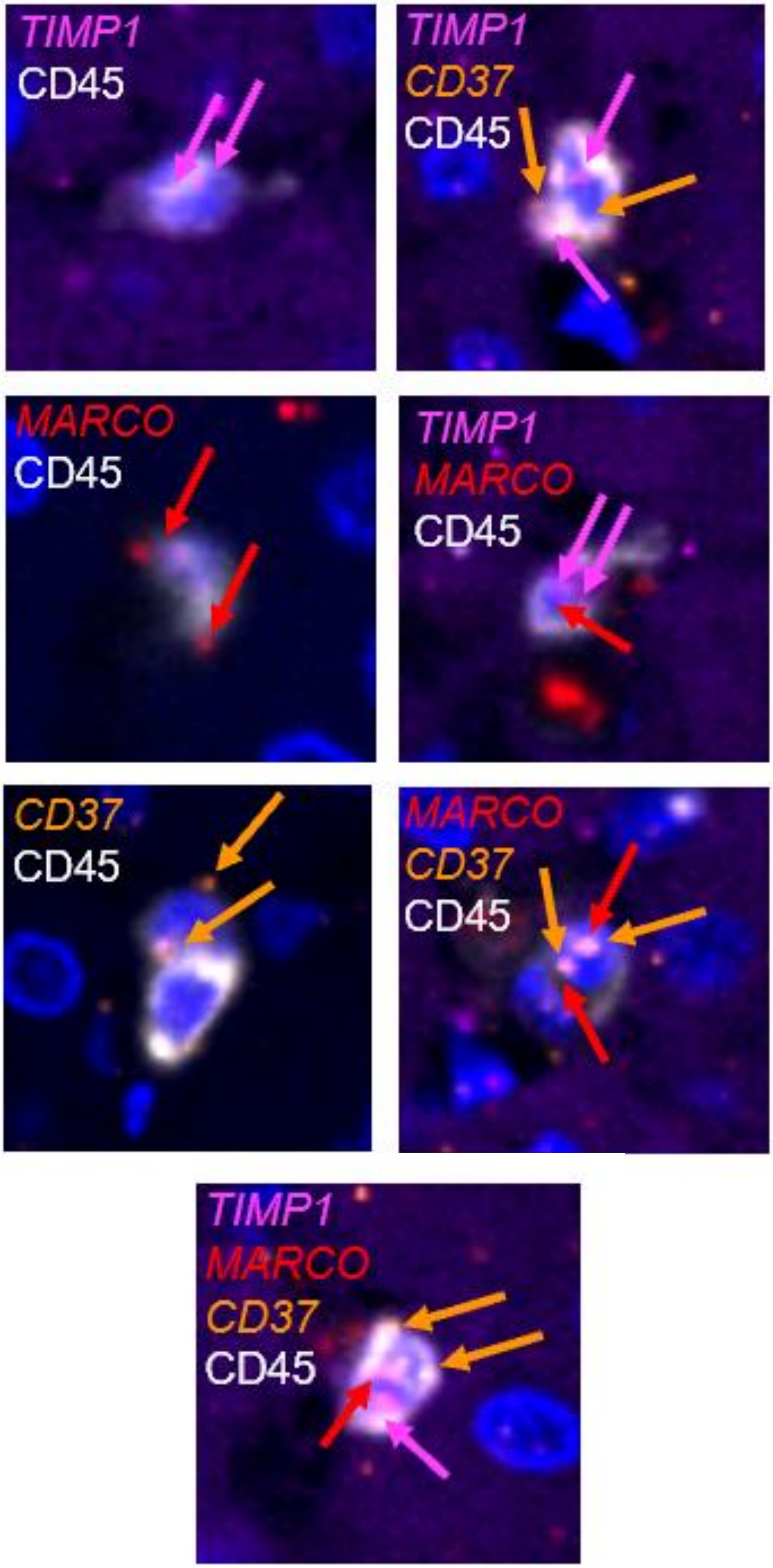


**Supplementary Fig. 5.** Representative images from multiplex RNAscope analysis of 3YO NHP livers. Images show autofluorescence to identify hepatic cords, DAPI-stained nuclei, CD45$^{+}$ immune cells identified by antibody, and RNA probe signals for *TIMP*, *MARCO*, *CD37*, and the overlay of all channels.

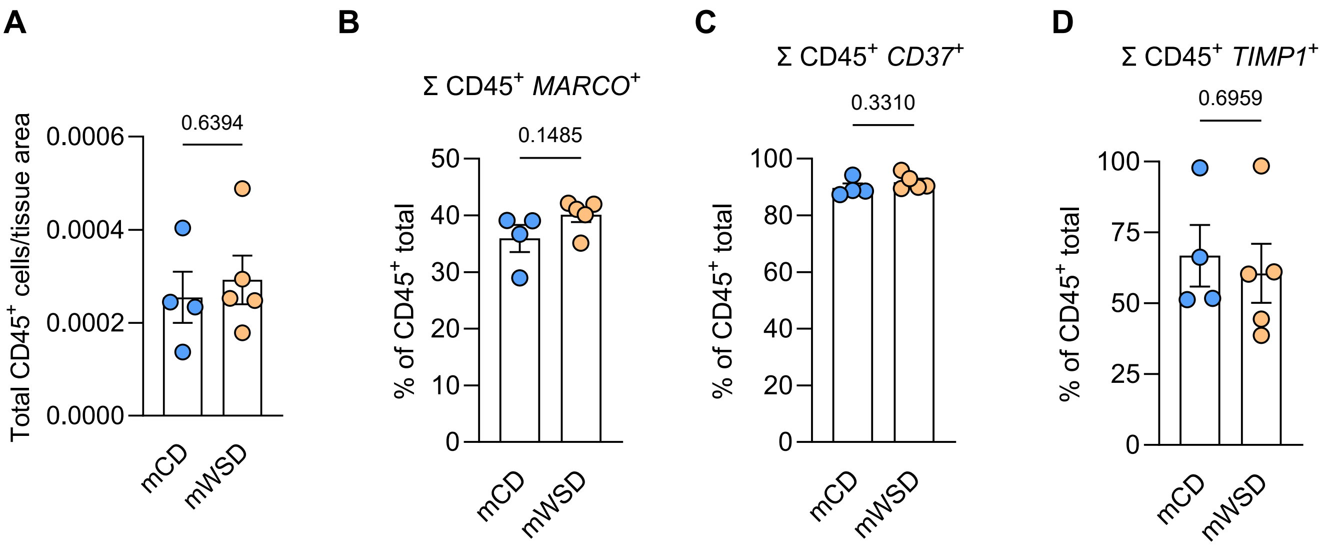


**Supplementary Fig. 6.** Co-expression of immune cell markers in 3YO NHP livers using RNAscope analysis. **A** Quantification of total CD45⁺ immune cells relative to total tissue area. Sum of all CD45⁺ immune cells co-expressing **B** *MARCO*, **C** *CD37*, and **D** *TIMP1*. Data are mean ± SEM. n = 4 mCD, n = 5 mWSD. Results were analyzed by Student's *t* test.

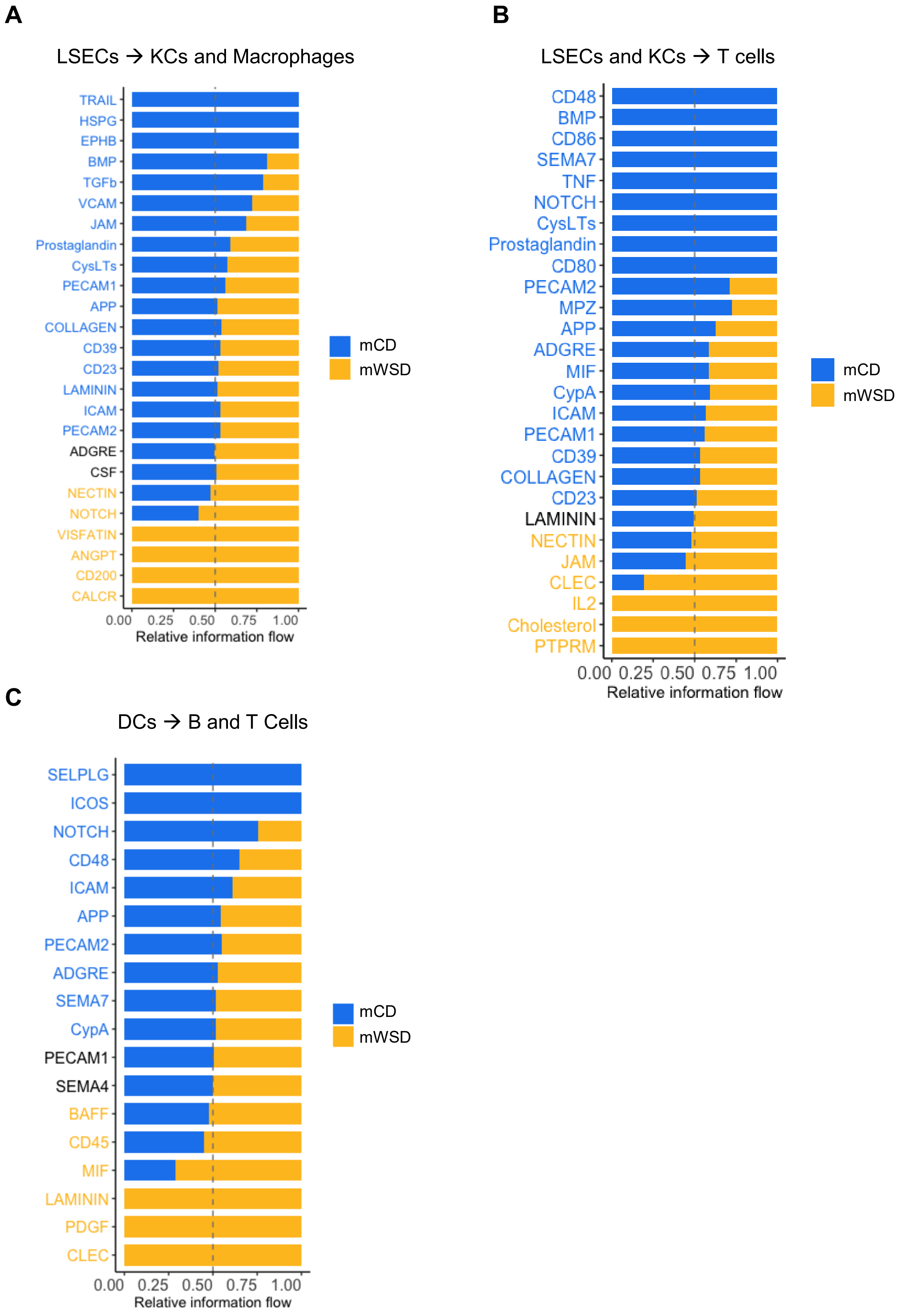


**Supplementary Fig. 7.** Pathways that are present or absent by maternal diet in (**A**) LSECs to KCs and macrophages, (**B**) LSECs and KCs to T cells, and (**C**) DCs to B and T cells. Blue or yellow pathway names indicate significant differences between maternal diet determined by Wilcoxon test (*P* < 0.05).
